# Analysis of developmental gene expression using smFISH and *in silico* staging of *C. elegans* embryos

**DOI:** 10.1101/2024.05.15.594414

**Authors:** Laura Breimann, Ella Bahry, Marwan Zouinkhi, Klim Kolyvanov, Lena Annika Street, Stephan Preibisch, Sevinç Ercan

## Abstract

Regulation of transcription during embryogenesis is key to development and differentiation. To study transcript expression throughout *Caenorhabditis elegans* embryogenesis at single-molecule resolution, we developed a high-throughput single-molecule fluorescence in situ hybridization (smFISH) method that relies on computational methods to developmentally stage embryos and quantify individual mRNA molecules in single embryos. We applied our system to *sdc-2*, a zygotically transcribed gene essential for hermaphrodite development and dosage compensation. We found that *sdc-2* is rapidly activated during early embryogenesis by increasing both the number of mRNAs produced per transcription site and the frequency of sites engaged in transcription. Knockdown of *sdc-2* and *dpy-27*, a subunit of the dosage compensation complex (DCC), increased the number of active transcription sites for the X chromosomal gene *dpy-23* but not the autosomal gene *mdh-1*, suggesting that the DCC reduces the frequency of *dpy-23* transcription. The temporal resolution from *in silico* staging of embryos showed that the deletion of a single DCC recruitment element near the *dpy-23* gene causes higher *dpy-23* mRNA expression after the start of dosage compensation, which could not be resolved using mRNAseq from mixed-stage embryos. In summary, we have established a computational approach to quantify temporal regulation of transcription throughout *C. elegans* embryogenesis and demonstrated its potential to provide new insights into developmental gene regulation.

## Introduction

Temporal and spatial regulation of transcription plays a crucial role in defining cellular identities and maintaining proper function in multicellular organisms. Studies using single-molecule fluorescent in-situ hybridization (smFISH) have quantified mRNA molecules and revealed that transcription is a highly stochastic process that occurs in discrete bursts over time [1–4]. Simple models of transcription include two key parameters: burst size, which describes the number of RNAs transcribed from a single promoter during an active period, and burst frequency, which quantifies the number of transcription bursts occurring over a given timeframe [1,5,6].

In embryogenesis, zygotic transcription controls early processes, including sex determination, dosage compensation, and body patterning [7]. In *Drosophila* embryos, developmental genes were shown to be subject to burst frequency modulation [2,8–10]. Studies including different organisms hypothesize that burst size is a property of promoters but was also shown to be controlled by transcription factors and chromatin regulatory mechanisms [11–17].

Similar to other organisms, transcriptional bursting has been observed in *Caenorhabditis elegans* [18,19]. However, the mechanisms that control the parameters of embryonic transcription have not been addressed. In *C. elegans*, zygotic transcription starts around the 4-cell stage [20–23]. Sex is determined shortly thereafter, and in XX animals, leads to zygotic transcription of *sdc-2*, which is essential for hermaphrodite development and X chromosome dosage compensation [24].

SDC-2 binds to the X chromosomes around the ∼40-cell stage and recruits the dosage compensation complex (DCC), which is composed of a specialized condensin I (hereafter DC) and several non-condensin proteins [25,26]. DCC represses X chromosome transcription ∼2-fold in XX hermaphrodites to equalize overall output from that of XO males [27,28]. Embryos defective for dosage compensation die late in embryogenesis, suggesting that this is a critical developmental time for proper gene dosage [29].

*C. elegans* is an excellent model for studying developmental gene regulation due to defined cell numbers and constant (stereotypic) lineage [30]. However, a major difficulty of performing smFISH at high throughput across embryogenesis is that there are no current methods to synchronize *C. elegans* embryos at specific stages. Accordingly, previous studies used smFISH in *C. elegans* to detect transcription qualitatively [31,32] with low throughput [19,33] or in early embryo stages [34,35].

To overcome the difficulty of obtaining smFISH measurements from distinct embryonic stages, we developed a computational approach to *in silico* stage embryos stained by smFISH probes and a nuclear marker (DAPI) acquired at random time points throughout embryogenesis. Our method suggests that the rapid zygotic activation of *sdc-2* is accomplished by increasing both the size and frequency of transcription bursts in early embryogenesis. We also quantified the effect of *sdc-2* and *dpy-27* knockdown on *dpy-23*, a dosage compensated X chromosomal gene, and used simultaneous staining for an autosomal gene *mdh-1* as an internal control. Our results show that the DCC represses *dpy-23* by reducing the frequency of active transcription sites per embryo. We compared the effect of a single DCC recruitment deletion using mRNA-seq from mixed-stage embryos versus our method and found that *in silico* staging coupled with high-throughput smFISH detects small temporal changes in gene expression during embryogenesis.

## Results

To obtain samples spanning all embryonic stages, we collected younger embryos by bleaching *C. elegans* adult hermaphrodites and older embryos by washing them off plates (Figure 1). We labeled single RNA molecules using smFISH probes targeting exonic or intronic regions of several genes on the X chromosome and autosomes (Supplemental Figure 1). We then semi-automatically imaged over 9000 smFISH-labeled embryos in 3D.

**Figure 1.**
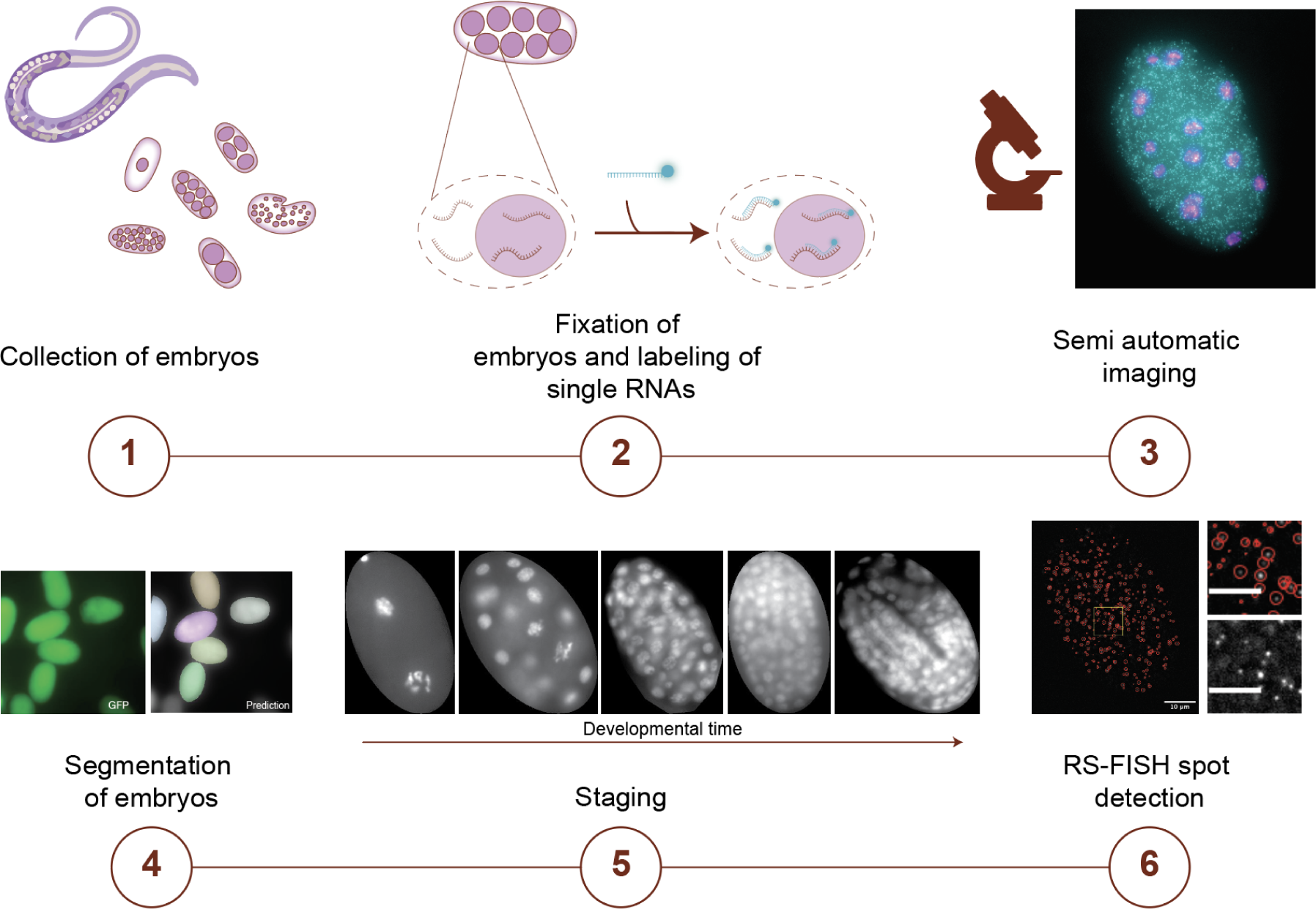
The experimental workflow. (1) Embryos are collected by bleaching adults and washing off older embryos. (2) Collected embryos are fixed, permeabilized, and labeled with smFISH probes. (3) The labeled embryos are imaged using a semi-automatic imaging setup. The analysis includes segmentation of single embryos (4), staging of embryos according to their developmental time (5), and detection of single RNAs using RS-FISH (6).

### Computational staging of *C. elegans* embryos and smFISH analysis

Imaging transcription using smFISH allows for additional visualization of nuclei by staining DNA with DAPI. Given that *C. elegans* follows a stereotypical developmental pattern, the number of nuclei directly correlates with its developmental stage [36,37]. While it is theoretically feasible to discern each embryonic stage ranging from a single nucleus to 558 nuclei at the conclusion of embryogenesis, such an undertaking would be exceedingly challenging for a high-throughput approach [38–40] or require staining of additional markers [41]. Therefore, we employ an autoencoder-based image classifier to learn representations from fixed, DAPI-stained *C. elegans* embryos to automatically stage them in an approach similar to [42]. We rely on a manually created training dataset for each developmental stage to achieve reliable predictions from the learned representations. Our training dataset was generated by categorizing embryos into different developmental stages.

The categories reflected the following stages of gene regulation and development: Bin 0 contains 1-4 cell stage embryos, which is before zygotic genome activation (ZGA) for most genes, thus detecting maternal mRNA load [21,22]. Bin 1 includes embryos with 5 - 30 nuclei, which start zygotic transcription, undergo sex determination, and begin to express SDC-2 [24,43]. Bin 2 includes embryos with 31-99 nuclei when DCC starts localizing to the X chromosomes [24,44,45]. Bin 3 includes embryos from 100 - 149 nuclei when H4K20me1 on the X chromosomes increases compared to autosomes [46–48]. Bin 4 includes embryos with nuclei from 150 - 534, where the nuclei are dense, and counting is especially difficult. In bin 5 (535 - 558), cells hardly divide, but embryos undergo several detectable morphological changes. We annotated at least 26 embryos per stage in the training set (for availability, see the methods section) (Figure 2B).

**Figure 2.**
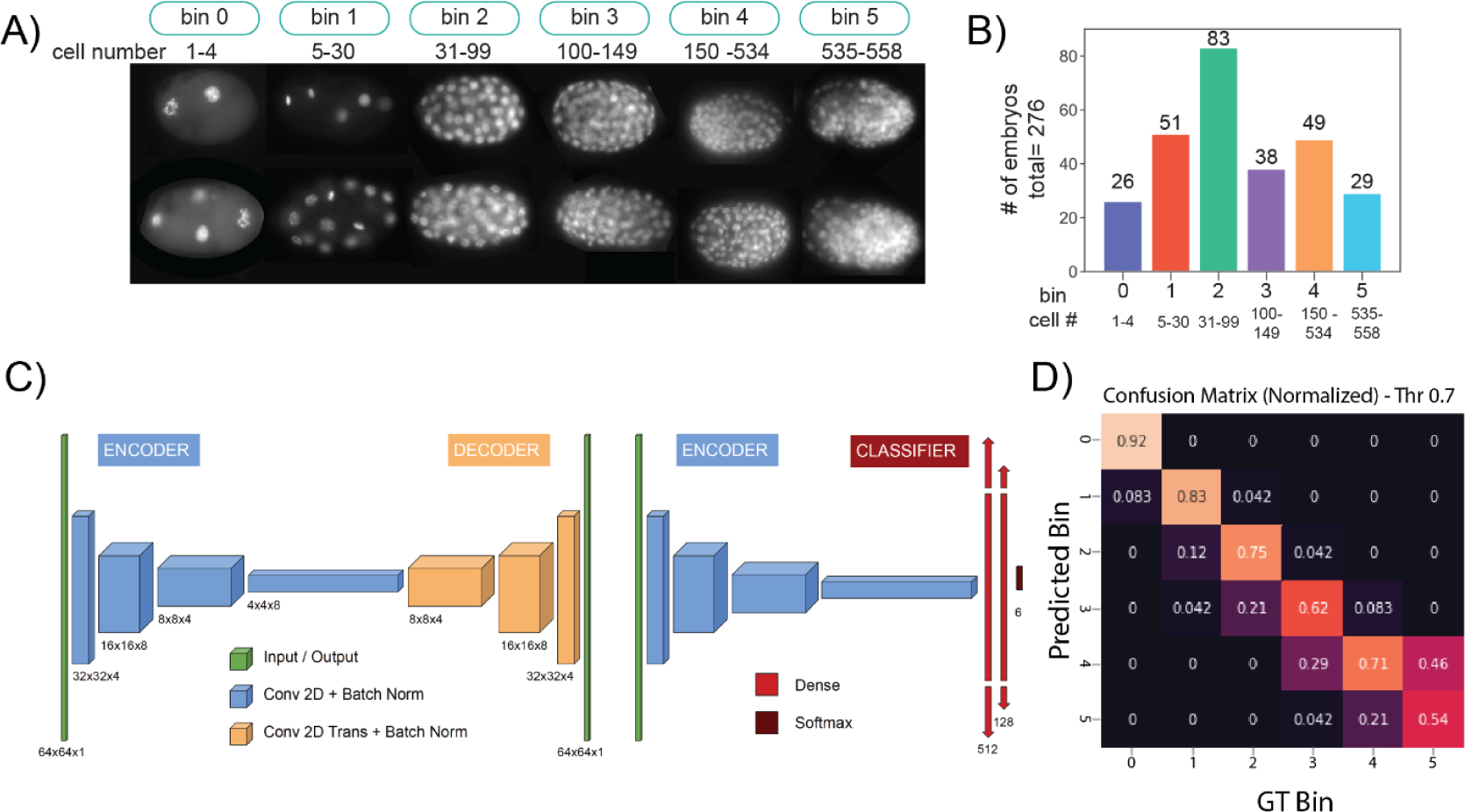
Prediction of embryonic stages using an autoencoder. A) Grouping all developmental stages into six bins with the included nucleus count labeled below each bin and additional example images of DAPI-stained embryos. **B)** Count of the annotated example images of each bin. The bin with the lowest count was used as a cut-off for all bins to have equal-sized training data. **C)** Illustration of the autoencoder architecture that was used for the classification of the stage bin. **D)** Confusion matrix for the predicted bins against a set of ground truth (GT) bins using a class-stratified leave-one-out validation. Values closer to 1 represent better predictions for the bin

For automatic stage prediction, we adapted an autoencoder-based architecture (Figure 2C) [49,50]. An autoencoder learns representative features of an input dataset, called the latent space, from which it tries to reconstruct the input dataset. Thus, it can be used as a non-linear dimensionality reduction method as the latent space captures the essential features that describe the data. First, to train the autoencoder, the entire dataset was utilized in a self-supervised manner, with the autoencoder learning to reconstruct the input data as its output, while 250 embryo images were withheld for validation. Next, we discarded the decoder and attached two fully connected layers directly to the encoder’s latent space output, forming an additional classifier arm designed to predict embryo stages. The classifier was initially trained by itself using the annotated training dataset, with the encoder weights frozen later, the entire network was trained end-to-end. Our approach provided sufficient prediction accuracy of the different embryonic stages (Figure 2D). Due to the dense packing of nuclei, early embryonic stages are more accurately predicted than older stages, which also holds true for the manual staging of embryos. While our approach has some uncertainty, especially for older embryonic stages, we decided that the ability to stage thousands of embryos automatically using only a small training set in a short time outweighs the benefits of the entirely (also uncertain) manual approach.

To quantify transcription in the *in silico* staged embryos, we implemented a pipeline for image analyses (Suppl. Fig. 2). Briefly, the raw smFISH signal was enhanced using a Median filter to reduce background noise. The individual transcripts were detected in 3D using the RS-FISH plugin [51]. The transcripts were filtered using a binary mask to exclude smFISH detections from neighboring embryos. Since imaging deeper into the tissue of embryos leads to loss of signal due to bleaching and scattering of light, we corrected the signal in Z by fitting a quadratic function to the detected spots. To compare transcripts from different embryos and conditions, we normalized the detections from each embryo by fitting a gamma function, where the maximum is set to 1.

### RNA Polymerase II knockdown causes distinct effects on the mRNA levels of different genes

To validate our analysis of transcription in embryos, we depleted the large subunit of RNA Pol II (AMA-1) by feeding bacteria expressing double-stranded RNAs for interference (RNAi). Since *ama-1* RNAi leads to germline formation defects, we used a relatively short exposure to allow embryo development. This treatment results in partial knockdown of mRNAs (Figure 3A).

**Figure 3.**
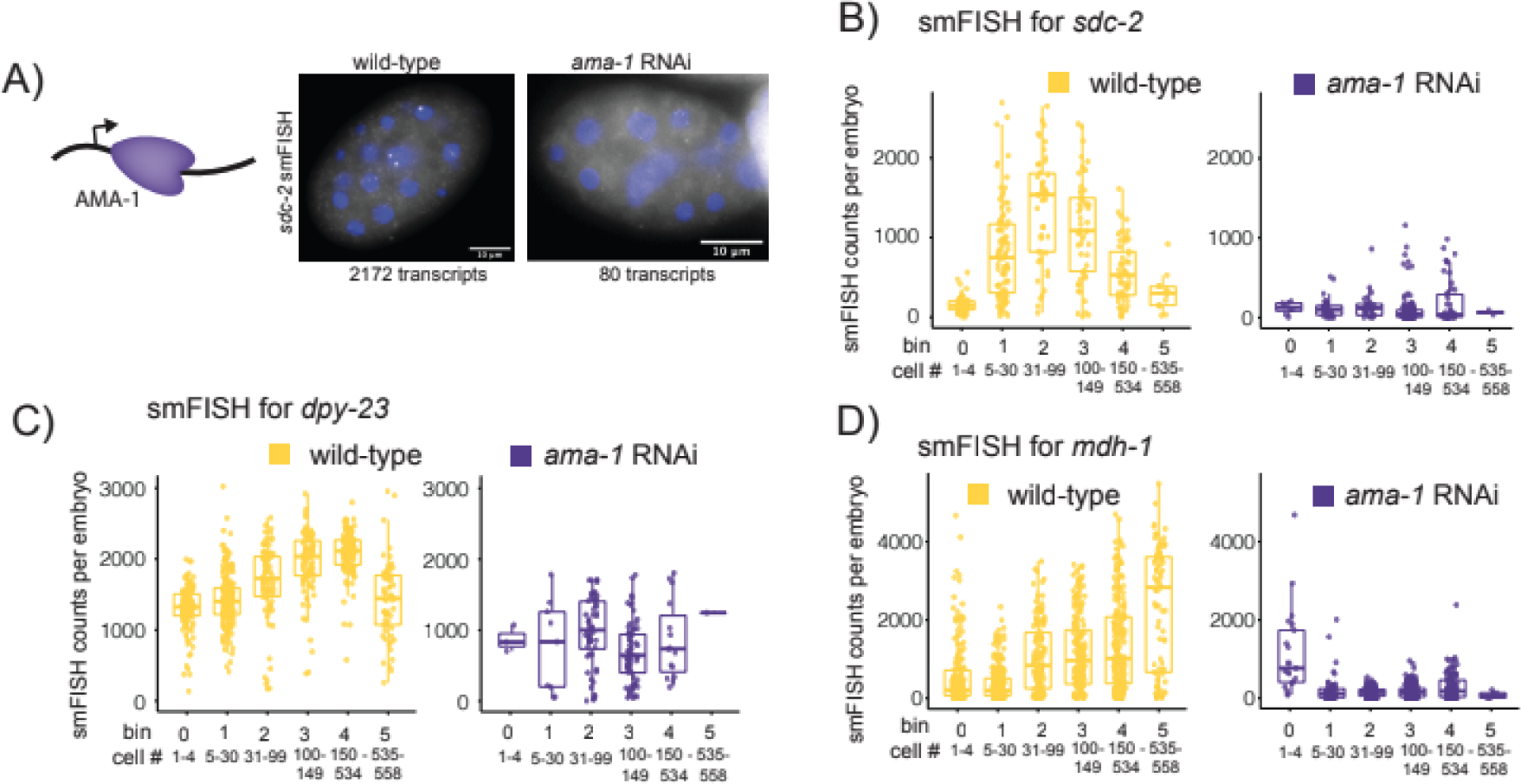
Detection of smFISH in wild-type and *ama-1* RNAi embryos. **A)** Cartoon of the RNA Pol II with the largest *C. elegans* subunit AMA-1 and example images of wild-type or *ama-1* RNAi embryos, labeled with *sdc-2* smFISH. The number of detected smFISH spots for each example is written under the image. **B)** Plots for smFISH detection of X chromosomal mRNA *sdc-2*. Each spot represents the total mRNA count for one embryo, plotted against the binned nuclei number for wild type (yellow) or *ama-1* RNAi (dark purple). **C)** Same as in B but for the X chromosomal gene *dpy-23*. **D)** Plot as in B but for the autosomal mRNA *mdh-1*.

The *sdc-2* gene is zygotically transcribed downstream of the sex determination pathway [24,43]. Accordingly, smFISH analysis shows few transcripts in the 1-4 cell stage (Figure 3B, left panel). *sdc-*2 is strongly activated at the 5-30 nuclei time-point, in agreement with its functions in early embryogenesis. *Ama-1* RNAi effectively prevents the activation of *sdc-2* transcription through embryogenesis (Figure 3B, right panel).

In contrast to the zygotically transcribed *sdc-2*, the maternally loaded *dpy-23* transcripts were reduced but were not eliminated by *ama-1* RNAi (Figure 3C). Interestingly, another maternally loaded mRNA, *mdh-1,* was present in 1-4 stage embryos but strongly reduced after the 4-cell stage, suggesting that the *mdh-1* mRNA has lower stability compared to *dpy-23* mRNA (Figure 3D). The distinct effects of *ama-1* knockdown on the three genes validate our methods and highlight the expression differences between *sdc-2*, *dpy-23,* and *mdh-1*.

### Distinguishing transcription sites from mRNAs using smFISH data

smFISH probes against exons detect single mRNAs as distinct spots with similar intensities [3,52]. The images also capture a few spots with high intensity inside the nucleus, marking sites of transcription that contain multiple mRNAs [4]. RNAi leads to the degradation of mRNA but does not affect *de novo* transcription. Accordingly, while *ama-1* RNAi reduced the high-intensity sites in *sdc-2* smFISH due to Pol-II reduction, *sdc-2* RNAi reduced overall mRNAs, while the high-intensity spots remained prominent (Figure 4A). Thus, our analysis supports the conclusion that high-intensity smFISH spots are sites of active transcription (Figure 4A).

**Figure 4.**
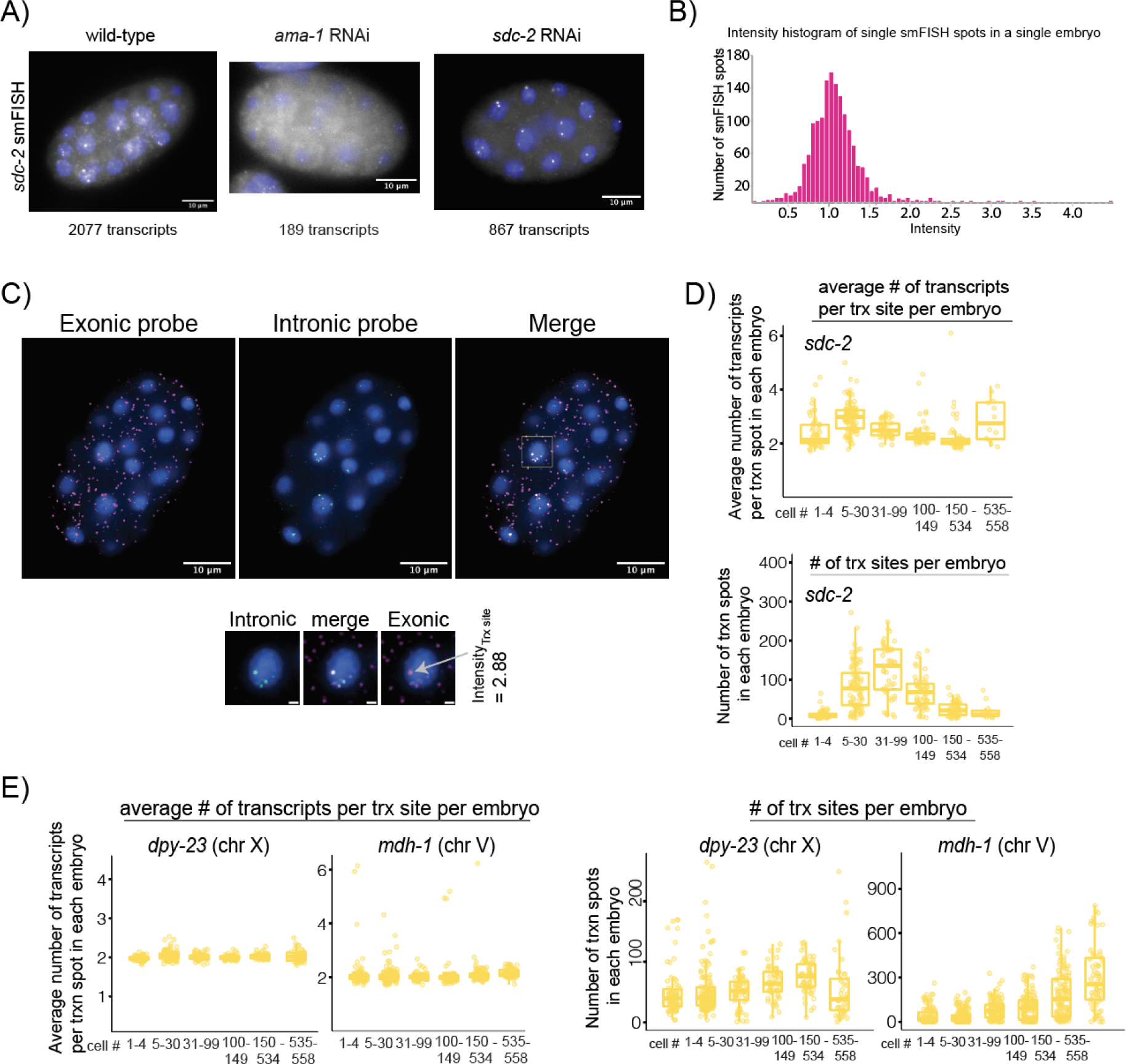
Detecting transcription sites and parameters. **A)** Example images of embryos labeled with *sdc-2* smFISH probes. Embryos are either wild-type, *ama-1* RNAi, or *sdc-2* RNAi. The number of detected transcripts is stated under each image. **B)** Histogram of normalized smFISH detections (exon) for *dpy-23* for a single embryo. **C)** Example embryo showing the dual labeling of *dpy-23* exonic (magenta) and intronic (green) smFISH probes. The lower panel shows a detailed view of the yellow boxed nucleus with the scale bar indicating 1 µm. The normalized intensity of the exonic transcription spot highlighted by the arrow is 2.88. The images were filtered using a Gaussian filter (sigma = 5). **D)** The upper plot shows the average number of *sdc-2* transcripts per transcription site per wild-type embryo. The lower plot shows the number of transcription sites per embryo. **E)** Left panel plots show the average number of *dpy-23* and *mdh-1* transcripts per transcription site in wild-type embryos—the plots on the right show the number of transcription sites per embryo.

To distinguish transcription sites from individual transcripts, first we normalized the smFISH signal by setting the maximum of the intensity distribution of all detected mRNAs per embryo to 1 (Figure 4B, Suppl. Fig. 2B), which corresponds to a single mRNA as previously established [4]. We used a threshold of 1.7 to capture spots with more than 1.7 full-length RNAs and interpret them as sites of active transcription. This approach might underestimate the number of active transcription sites as genes that just started being transcribed would be excluded. To validate our approach, we used smFISH probes against introns, which mark the site of active transcription [4,52]. Visual inspection of several co-stained transcription sites confirmed this threshold (Figure 4C). Occasionally, more than two intronic spots were observed inside or outside a nucleus, possibly due to intron retention [53,54]. Since exonic probes allow the normalization of smFISH intensity between embryos, we used the exonic smFISH data to analyze transcription sites.

### Zygotic activation of *sdc-2* involves regulation of both transcription burst size and frequency

In the two-state random telegraph model of transcription kinetics, an increase in transcription can be accomplished by increasing the burst frequency (how often a promoter is turned on) or the burst size (the number of RNAs produced at a transcription site) [55,56]. These modeling approaches use experimental measurements in mammalian systems that are currently lacking for *C. elegans* [57,58]. Therefore, we reasoned that the number of transcription sites detected in each embryo (normalized to the number of nuclei) correlates with the frequency of transcription bursts, and the smFISH intensity detected at each transcription site correlates with the size of the transcription bursts.

Analysis of *sdc-2* in wild-type embryos showed that both the number of transcripts per transcription site and the number of transcription sites per embryo increase in the 5-30 cell stage when *sdc-2* is transcription activated for functions in sex determination and dosage compensation (Figure 4D). For *dpy-23* and *mdh-1*, the intensity of the smFISH signal at transcription sites was more similar, while the number of sites varied across embryogenesis (Figure 4E).

### Knockdown of the DCC subunits SDC-2 and DPY-27 leads to increased transcription of *dpy-23*, a dosage-compensated gene on the X chromosome

The DCC subunit SDC-2 is the first to bind the X chromosomes and is required for the recruitment of the rest of the complex (Figure 5A) [24,59]. In hermaphrodite embryos, the knockdown of SDC-2 results in a ∼1.8-fold increase of RNA Pol II binding at X chromosomal promoters as measured by GRO-seq [27]. We knocked down SDC-2 using RNAi and analyzed the transcription of a dosage-compensated gene, *dpy-23*. We used mock RNAi as a control because feeding is performed on different growth media and uses a different strain of bacteria compared to standard growth conditions used for the wild-type strain N2. We validated the level of *sdc-2* knockdown (Suppl. Fig. 3A) by quantifying the *sdc-2* smFISH signal (Suppl. Fig. 3B).

**Figure 5.**
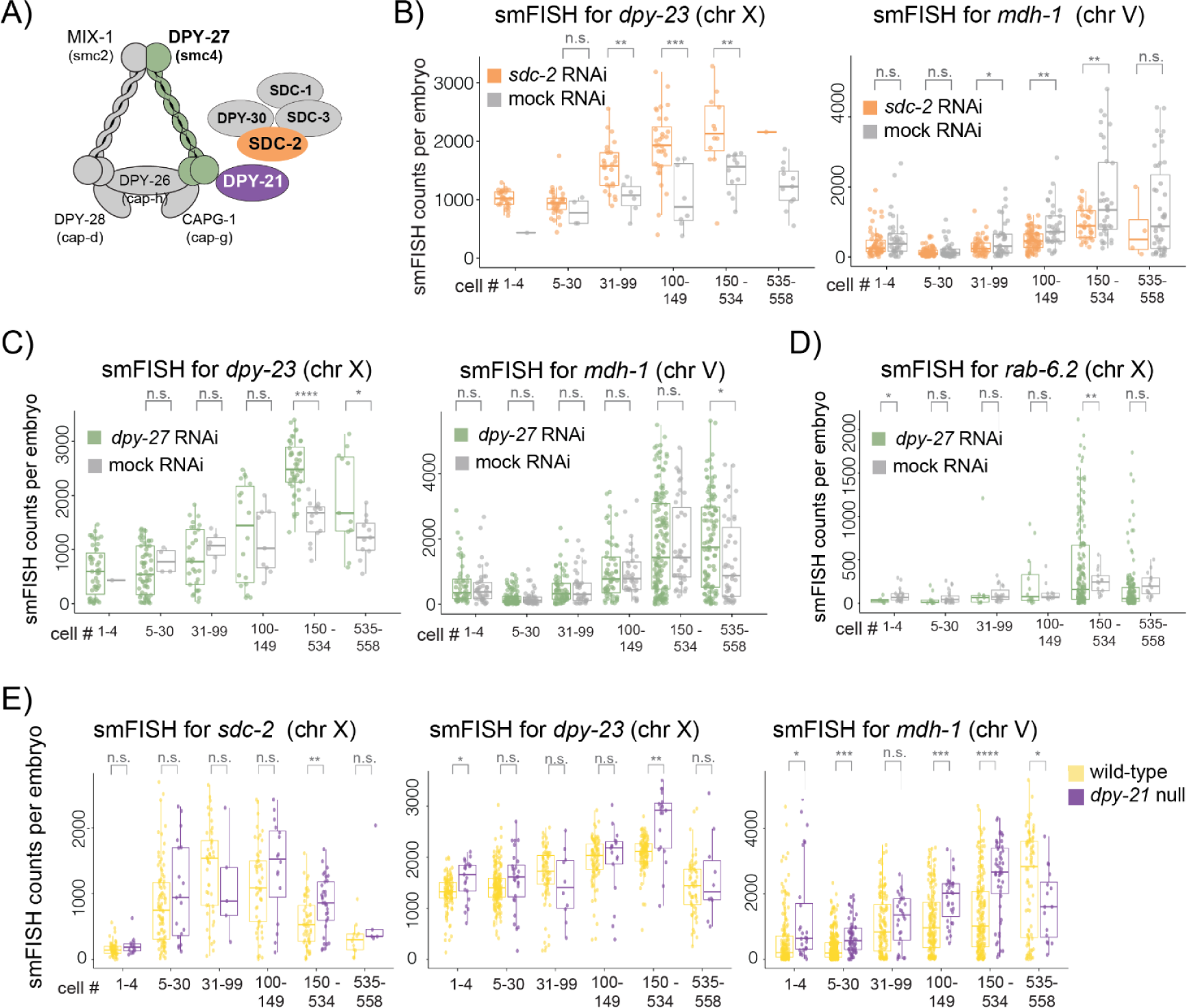
Measuring the effect of DCC knockdown on a dosage-compensated gene. **A)** Cartoon of the DCC complex highlighting the subunits SDC-2, DPY-27, and DPY-21. **B)** Plots for smFISH detection of X chromosomal mRNA *dpy-23* and autosomal mRNA *mdh-1*. Each spot represents the total mRNA count for one embryo, plotted against the binned nuclei number for wild type (yellow) and *sdc-2* RNAi (orange) Data plotted for *mdh-1* includes measurements from additional embryos that were probed for *sdc-2*. **C)** Plots for the X chromosomal mRNA *dpy-23* or the autosomal RNA *mdh-1* in mock RNAi treatment (gray) or *dpy-27* KD (green). Data plotted for *mdh-1* includes embryos probed for *dpy-23* and *rab-6.2*. **D)** Plots for the X chromosomal gene *rab-6.2* in mock RNAi (gray) or *dpy-27* KD (green) condition. **E)** Plots for the X chromosomal genes *sdc-2* and *dpy-23* as well as the autosomal gene *mdh-1* in wild-type (yellow) or *dpy-21* KO (purple) conditions. Data plotted for *mdh-1* includes embryos probed for *sdc-2* and *dpy-23*. Significant P-values from Welch’s unequal variances t-test are noted above the wild-type vs. knock-down condition for each time-point. P-values are denoted as n.s. (P > 0.05), * (P ≤ 0.05), ** (P ≤ 0.01), *** (P ≤ 0.001) and **** (P ≤ 0.0001). If one of the samples had less than two points, no P-value was calculated.

To capture the biological and technical variability between embryos, transcripts from an autosomal gene *mdh-1* were measured in the identical embryos in parallel to *dpy-2*3. Since RNAi produces variable knockdown, having a large number of embryos was important to test for statistical significance. Where the sample numbers were low, we refrained from performing a statistical test. We found that *sdc-2* knockdown leads to upregulation of *dpy-23* but not *mdh-1* (Figure 5B). Notably, upregulation of *dpy-23* mRNA started from the 31-99 cell stage, immediately following DCC localization but before H4K20me1 enrichment on the X chromosomes. The apparent downregulation of *mdh-1* upon *sdc-2* RNAi may be due to the previously demonstrated repressive effect of *sdc-2* knockdown on autosomes [60].

To validate that the upregulation of *dpy-23* was due to reduced DCC binding, we performed RNAi to knock down the condensin core of the DCC, targeting the *smc-4* variant *dpy-27* (Suppl. Fig. 3C, D). *Dpy-27* RNAi also resulted in an increase in *dpy-23* transcripts (Figure 5C). To test if DCC-mediated repression can be observed for another X chromosomal gene whose transcription profile differs from that of *dpy-23*, we analyzed *rab-6.2*, a dosage-compensated gene with a lower transcription level and no maternal loading. There was higher variability in *rab-6.2* transcription in the *dpy-27* RNAi condition (Figure 5D). The reason for this variability is unclear.

### Analysis of transcripts in a null mutant of DPY-21 H4K20me2 demethylase

Condensin DC recruits the H4K20me2 demethylase DPY-21 to the X chromosomes, resulting in the enrichment of H4K20me1 starting at ∼100-cell stage [46,48,61,62]. DPY-21 loss is not lethal thus, a homozygous null line is viable and shows a pronounced dumpy phenotype due to increased X chromosomal gene expression [46,59,62]. In the *dpy-21* null mutant embryos, the transcripts from two X chromosomal genes, *sdc-2* and *dpy-23,* increased in 150-534 nuclei (Figure 5E). Unlike other DCC subunits, DPY-21 is expressed in the germline and regulates H4K20me1 on the autosomes [62]. We observed an upregulation of *mdh-1* in the *dpy-21* null mutant (Figure 5E), which may reflect the repressive function of H4K20me1 on autosomal transcription.

### smFISH combined with *in silico* staging detects the local and temporal effect of an individual DCC recruitment site deletion

In embryos, DCC binding across the length of the ∼17.7 Mb X chromosome initiates at a set of ∼60 recruitment elements on the X (*rex*) [26]. The DCC spreads out from these sites and exerts its repression chromosome-wide [28,63]. Individual *rex* sites collectively contribute to recruitment and repression, thus deletion of one or few *rex* sites causes subtle changes, as measured by mRNA-seq in mixed-stage embryos [64,65]. We wondered if our approach could reveal the contribution of an individual *rex* site in developmentally staged embryos.

We previously constructed a strain deleting *rex-1*, located ∼6 kb downstream of *dpy-23* [64]. Using three different fluorophores (Suppl. Fig. 1B), we quantified two internal control genes simultaneously with *dpy-23*; *wdr-5.2,* which is far from *rex-1,* and the autosomal gene *mdh-1* (Figure 6A). Compared to the two internal controls, *dpy-23* transcripts increased significantly upon *rex-1* deletion (Figure 6B, Suppl. Fig. 5).

**Figure 6.**
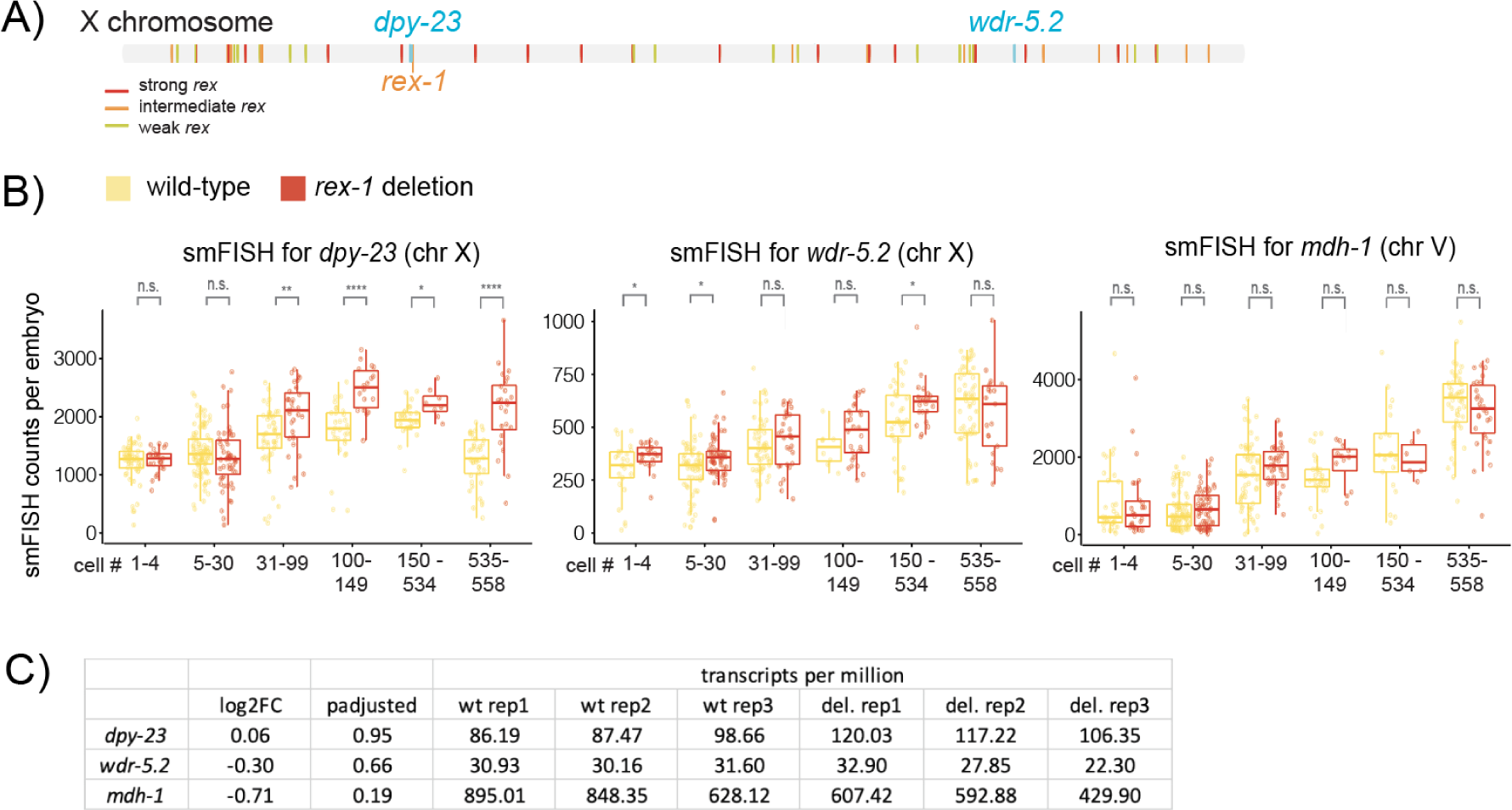
Influence of a single rex site deletion on a neighboring gene. **A)** Cartoon of the X chromosome with approximate locations of the genes *dpy-23* and *wdr-5.2* (∼10 Mbp away) and the location of *rex-1*. **b)** Plots for smFISH detection X chromosomal mRNAs (*dpy-23* and *wdr-5.2*) and one autosomal mRNA (*mdh-1*) in wild-type and *rex-1* deletion strains. The data are from the same embryos using 3-color imaging. Each spot represents the total mRNA count for one embryo, plotted against the binned nuclei number for wild-type (yellow) or *rex-1* deletion (red) strains. Significant P-values from Welch’s unequal variances t-test are noted above the wild-type vs. deletion condition for each time point. P-values are denoted as n.s. (P > 0.05), * (P ≤ 0.05), ** (P ≤ 0.01), *** (P ≤ 0.001) and **** (P ≤ 0.0001). **c)** mRNA-seq data from mixed-stage embryos isolated from gravid adults was used for differential expression analysis. DESeq2 output of the log2 fold change and adjusted p values are provided along with the expression values from the three biological replicates.

Next, we tested whether mRNA-seq could detect the same effect in mixed-stage embryos. The transcript per million (TPM) measure of *dpy-23* expression was slightly elevated in the deletion strain but did not reach statistical significance (Figure 6C). This is likely due to mixed-stage embryos containing a high proportion of early embryos that have not started dosage compensation. Thus, smFISH analysis of *in silico* staged embryos reveals the effect of a *rex* site that is missed by population assays.

### SDC-2 and DPY-27 knockdown increases the number of *dpy-23* transcription sites per embryo

Next, we addressed how the DCC represses *dpy-23* expression. Upon *sdc-2* RNAi, the average *dpy-23* smFISH signal at the transcription sites did not change, but the number of transcription sites per embryo increased significantly (Figure 7A). The effect of *dpy-27* RNAi on *dpy-23* was similar to that of *sdc-2*, increasing the number of transcription sites per embryo rather than the number of transcripts at each site (Figure 7B). To ensure that our conclusion is not sensitive to threshold selection with which we define active transcription sites, we tested if an intensity threshold of 1. 5 or 2.0 changes the result. Both thresholds recapitulated the main finding (Suppl. Fig. 4). In addition to the RNAi condition, in the *rex-1* deletion strain, the number of transcription sites increased significantly for the neighboring gene *dpy-23* (Figure 7C) and not for the control genes (Suppl. Fig. 4). Thus, our results support the idea that the DCC represses *dpy-23* transcription by reducing its frequency of transcription.

**Figure 7.**
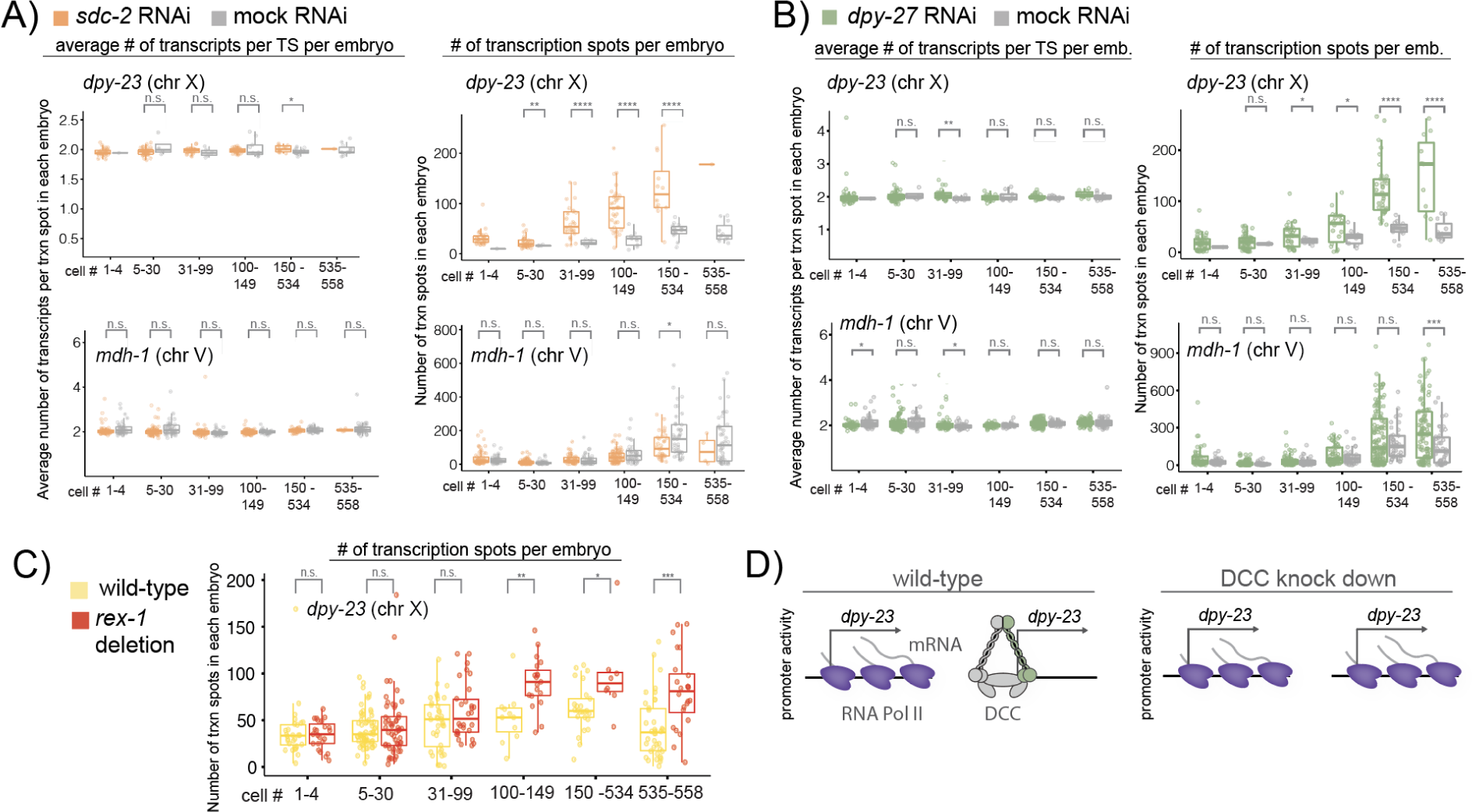
Analysis of transcription site parameters upon DCC knockdown. **A)** Relative transcription parameter (average number of transcripts per transcription site per embryo and the number of transcription spots per embryo) changes for the X chromosomal gene *dpy-23* for *sdc-2* RNAi (orange) and mock RNAi (gray) conditions. The left plots show the average number of nascent transcripts per TSS, which is correlated with the burst size. The right plots show the number of actively transcribing promoters, which is correlated with the burst frequency. Data from *mdh-1* includes embryos co-probed with *dpy-23* and *rab-6.2*. **B)** Relative transcription parameter changes for *dpy-27* RNAi (green) and mock RNAi (gray) conditions. **C)** Relative number of transcription spots per embryo for the X chromosomal gene *dpy-23* for wild-type (yellow) and *rex-1* deletion (red) conditions. Significant P-values from Welch’s unequal variances t-test are noted above the wild-type vs. knock-down condition for each time-point. P-values are denoted as n.s. (P > 0.05), * (P ≤ 0.05), ** (P ≤ 0.01), *** (P ≤ 0.001) and **** (P ≤ 0.0001). If one of the samples had less than two too few points, no P-value was calculated. **D)** Model for transcription regulation during dosage compensation.

## Discussion

smFISH analysis of transcription in developing organisms is powerful but presents experimental and computational challenges. In this work, we used widefield smFISH to quantify transcripts in each embryo and applied machine learning to stage a large number of embryos for analysis of transcription throughout embryogenesis in *C. elegans*.

### Transcriptional activation of an early zygotic gene sdc-2

Our analysis of *sdc-2* suggests that both the rate and frequency of transcription bursts increase significantly following zygotic genome activation (Figure 4). In *C. elegans*, the single fertilized cell undergoes several divisions to become a 30-cell-stage embryo in ∼150 min [36,66,67]. Rapidly activated zygotic genes tend to be shorter [53]. Yet, *sdc-2* transcript (∼10 kb) is longer than the median transcript in the genome (∼2 kb) and encodes a large ∼340 kD protein. Therefore, multiple mechanisms may activate *sdc-2* transcription to ensure robust production of SDC-2 protein for timely initiation of hermaphrodite development and dosage compensation.

### Timing of DCC-mediated transcription repression

Our current model is that the DCC starts repression soon after SDC-2-mediated recruitment of condensin DC to the X chromosomes and is augmented by enrichment of H4K20me1 in later embryogenesis [59,61]. Consistent with this model, we observed an earlier and stronger effect of *sdc-2* RNAi compared to *dpy-27* and *dpy-21* (Figure 5). The DCC-mediated repression starting after the 30-cell stage was also supported by the effect of *rex-1* deletion on the neighboring *dpy-23* gene (Figure 6). The deletion analysis also highlighted the power of *in silico* staging embryos to detect the temporal effect of a single *rex* site, which was missed by mRNA-seq analysis of mixed-stage embryos.

### Mechanism of DCC-mediated transcription repression

Our data suggest that the DCC reduces the number of *dpy-23* alleles that are actively transcribed in an embryo (Figure 7). Notably, *dpy-23* expression during development appears to be regulated by the frequency of transcription rather than the number of mRNAs being produced at each site (Figure 4E). Therefore, the DCC may repress *dpy-23* by modulating the mechanism that controls its frequency of transcription. It will be interesting to find another X chromosomal gene that is primarily activated by burst size to test if the DCC also has the ability to also modulate the size of transcription bursts.

How does the DCC reduce the frequency of *dpy-23* transcription? Previous research has shown that in addition to TFs [3,13–15,68–71] and chromatin [16,17,59,72–75], 3D organization of enhancers and promoters can modulate burst frequency [9–11,72,76–78]. The DCC has the potential to regulate one or more of these processes, as the core of the DCC is a structural maintenance of chromosomes (SMC) complex that controls the 3D DNA contacts on the X chromosomes [79,80] and is required for the compaction and regulation of histone modifications on chromatin [46,48,61,81–83].

The level and dynamics of condensin DC binding are important for transcription repression [59,81]. In a *dpy-21* null mutant where X chromosome transcription increases, the proportion of mobile DPY-27 proteins, as measured by fluorescence recovery after photobleaching (FRAP), decreases dramatically [59]. Therefore, we speculate that the transient association of condensin DC molecules with the *dpy-23* promoter may reduce RNA Pol II initiation at a fraction of the time, resulting in fewer alleles transcribing within a population of embryos (Figure 7D).

### Fine-tuning the transcriptional output of an entire chromosome for dosage compensation

Transcription is a stochastic process leading to differential mRNA and protein levels between cells and organisms with the same genetic background and sharing the same environment [84,85]. Even in a deterministic organism like *C. elegans* with a fixed lineage and cell numbers, transcription was described as stochastic in reaction to mutations [19] during development [18]. The presence of X chromosome dosage compensation mechanisms argues that the high level of transcriptional heterogeneity at the short time scale is averaged over time to produce the required dosage of gene transcripts.

In *D. melanogaster*, a complex of proteins and non-coding RNAs called the male-specific lethal (MSL) complex binds to and increases transcription from the X chromosome in males to accomplish dosage compensation [86,87]. A recent study used the MS2-MCP system to live-image transcription of four X chromosomal genes in female and male embryos and found that the amplitude of transcriptional bursts was increased in males [88]. How the different mechanisms of *D. melanogaster* and *C. elegans* dosage compensation (2-fold upregulation in the fly and 2-fold downregulation in the worm) modulate the burst and frequency parameters of transcription to fine-tune the average output of an entire chromosome is a fascinating question for future research.

## Methods

### Strains and Worm Growth

Unless noted, worms were grown and maintained using standard methods at 20-22°C on NGM plates (1L: 980 ml Milli-Q water, 3 g NaCl, 20 g agar, 2.5 g peptone, 1 ml 1 M CaCl2, 1 ml 5 mg/ml cholesterol in ethanol, 1 ml 1 M MgSO4 and 25 ml 1 M KPO4 buffer) containing OP50-1 strain of *E. coli* as food. Worm strains used in this study include N2 (wild-type, CGC), CB428 (dpy-21 (e428) V, CGC), and ERC41 (ers8[delX:4394846-4396180]) [64].

### RNAi conditions

RNAi conditions used in this study include pop-1 (W10C8.2), sdc-2 (C35C5.1), dpy-27 (R13G10.1), ama-1 (F36A4.7), rluc (Renilla luciferase) and empty vector (L4440) all from the Ahringer library. Bacteria were sequenced before use to ensure the correct strain was used. RNAi plates (RNAi plates (1 L), 3 g NaCl, 17 g bacto agar, 3 g bacto peptone, 975 ml water, autoclave, cool and add: 1 ml cholesterol (5 mg/ml in EtOH), 1 ml 1M CaCl_2,_ 1 ml 1M MgSO_4_, 25 ml 1 M K phosphate buffer, 1 ml IPTG (1 mM final concentration, made freshly), 1 ml Amp (50 mg/ml stock), 1 ml Tet (12 mg/ml stock)) were prepared freshly and usually not stored longer than one month in the dark. Single clones of bacteria were picked and cultured in a 5 ml overnight culture (LB + Amp Kan) at 37°C and shaking at 400 rpm. The next day, the small culture was added to typically 200 mL LB (Amp Kan) and continued shaking for 2-4 hours when the density of the culture reached OD 1. Then, 1 ml IPTG was added to the culture, and incubation continued for 2 hours. To reduce the bacteria volume by 1/10, bacteria were spun down at 1000 g and resuspended in M9 to have about 400 µl of bacteria per plate (very dense).

Worms were grown to the adult stage and bleached to retrieve embryos. Embryos were placed in a 15 ml tube with M9 at RT and were shaken overnight. The next day, starved L1s were placed on the freshly prepared RNAi plates at 20°C in the dark. For *ama-1* RNAi, L3/L4 were placed on RNAi bacteria to allow germline formation. After three days, adult worms were collected, and embryos were extracted, as described below.

### smFISH staining in *C. elegans* embryos

The smFISH labeling of *C. elegans* embryos was performed as reported in Bahry et al. [51] with some modifications.

### Probe design

All smFISH probe sets (Custom Stellaris® RNA FISH Probe Set) were designed and manufactured by Biosearch Technologies using the Stellaris probe designer web tool (Biosearch Technologies) specifically for *C. elegans*. Probes were designed to target exons or introns and labeled Quasar 670, CAL Fluor Red 610, or Quasar 570. Probes were designed to have at least two-nucleotide space in between neighboring probes to avoid quenching. For exonic or intronic probe design, sequences listed at wormbase.org were used, and splice regions were masked with “n” nucleotides to avoid probes in these areas. If possible, the maximum amount of probes (48 per order) was selected for each exonic or intronic sequence. An example of the distribution of exonic and intronic probes for *dpy-23* mRNA can be found in Supplemental Figure 1a. Filter sets with narrow bandwidths were selected to separate the detection from the three wavelengths (λ 570 nm, 610 nm, and 670 nm). We observed no bleedthrough between the three channels (Supplemental Figure 1b). Typically, far-red dyes performed better due to the autofluorescence of *C. elegans* embryos, especially in older stages. The number and color of probes are listed in Supplemental Table 1. The list of complete sequences can be found in the Supplements.

### Preparation of worms

Worms were synchronized by either bleaching or egg-laying and grown to the adult stage. At this step, having non-starved, healthy adult worms in sufficient amounts is crucial. Typically, at least 5 x 6 cm plates full of worms were used to collect embryos.

### Collection of embryos

For embryo collection, worms, and older embryos were washed off plates using M9 (5.8 g Na2HPO4, 3.0 g KH2PO4, 0.5 g NaCl, 1.0 g NH4Cl, Nuclease-free water to a final volume of 1000 ml) and collected in a 35 µm nylon filter. Worms were washed at least three times with H_2_O, then carefully transferred to a falcon tube using M9 and allowed to settle down. The supernatant was removed, and 5 ml of freshly prepared bleaching solution (2.5 ml 4N NaOH, 2.5 ml 5% NaClO, 5 ml Nuclease-free water) was added to the worms. The dissolving of the adult worms was closely observed, and after 3 minutes, the tubes were spun at 3000 g for 1 minute to collect the embryos. The supernatant was quickly removed, and the embryo pellet was vortexed. Then 10 ml of 1x PBS (with 0.05% of Triton X-100) was added, and the tube was centrifuged again for 3 min at 3000 g. This step was repeated twice until a clean embryo extract was left in the tube.

### Fixing and permeabilization

Embryos were resuspended in 1 ml fixation solution (4% paraformaldehyde (PFA) in 1xPBS (DEPC treated + autoclave + 0,05% of Triton X-100) and incubated at RT for 15 min while rotating. Next, the tube was submerged in liquid nitrogen for 1 minute to freeze crack the embryo eggshells. The tube was then transferred to a beaker with RT water to thaw, and once it was fully thawed, it was kept on ice for an additional 20 minutes. After this incubation, the tubes were spun down at 3000 g for 3 minutes, the supernatant was removed, and the embryos were washed twice with 1 ml of 1x PBS (with 0.05% Triton X-100). The embryos were resuspended in 70% EtOH and kept at 4 °C for at least 24 hours. Embryos can be kept at 4 °C for at least several weeks.

### smFISH staining

For smFISH staining of the previously fixed embryos, tubes were centrifuged at 3000 g for 3 minutes, and ethanol was carefully removed. The pellet can be loose at this step, so removing the supernatant should also be done in two stages. Embryos were then resuspended in 1 ml wash buffer (40 ml nuclease-free water, 5 ml deionized formamide, 5 ml 20x SSC) and vortexed. Tubes were centrifuged, as above, and the supernatant was removed. The embryos were then resuspended in 50 µl hybridization solution (50 µl H20 (RNAse free), 37.5 µl ethylene carbonate (EC) 5 mg/ml, 25 µl formamide (at RT), 12.5 µl SDS (dissolved), 125 µl dextran sulfate 10%), and 1 µl of each probe set (12.5 µM stock solution) was added directly to the sample. Tubes were then vortexed lightly and incubated at 37°C in the dark overnight. The next day, 0.5 ml of the wash buffer was added, and the tubes vortexed and centrifuged to remove the supernatant. Next, 1 ml of wash buffer was added, and samples were incubated at 37°C for 30 minutes. After that, tubes were centrifuged again to remove the supernatant, and the embryo pellet was vortexed before adding 1 ml wash buffer. In this step, DAPI (5 ng/mL) was added to the wash buffer, and tubes were incubated at 37 °C for 30 minutes. After centrifugation, the wash buffer was removed, and samples were washed once with 2x SSC.

### Mounting

Most liquid was removed from the stained embryos to mount the stained embryo tubes. About 15 µl of dense embryo solution (in 2x SSC) were used per microscopy slide and spread onto a coverslip (#1.5, 22 x 22 mm). The sample was left to dry for about 15 minutes, and then 15 µl of ProLong™ Diamond Antifade Mountant (Thermo Fisher) was added to the sample. A glass slide (SuperFrost, VWR) was pressed onto the embryos, and mounting media was placed on the coverslip. Slides were left at RT in the dark for 24 hours to cure before sealing the sides with nail polish and then stored at 4°C. Images were then acquired within two weeks of the sample being prepared.

### Imaging smFISH embryos

Embryos were imaged on a Nikon Ti inverted fluorescence microscope with an EMCCD camera (ANDOR, DU iXON Ultra 888), Lumen 200 Fluorescence Illumination Systems (Prior Scientific), and a 100x plan apo oil objective (NA 1.4) using appropriate filter sets (Gold FISH (Chroma 49304), Red FISH (Chroma 49306), Cy5 Narrow (Chroma 49396), GFP, DAPI). Images were acquired with 90 z-stacks positions with 200 nm step-width using Nikon Elements software. Every field of view had 1024x1024 pixels (XY axes) and ∼ 90 slices (Z), with 0.13 μm lateral and 0.2 μm axial resolution. Positions of embryos were marked in the software for semi-automatic imaging.

The 7447 images used in this study are provided at https://data.janelia.org/3Mq4F. The same site also provides 2505 smFISH images corresponding to additional conditions not used in this paper.

### Embryo instance segmentation

Single fields of view contained several embryos, which were segmented for downstream analysis. An instance segmentation was performed on the max projection of the GFP channel, which contained an autofluorescence signal marking the entire embryo (Supplemental Figure 2a). We used a 2D StarDist implementation, an approach for nuclei/cell detection with a star convex shape, to extract individual embryos [89]. A ground-truth dataset was created using a random forest classifier and a custom ellipsoid fit script implemented in Java, resulting in 1734 fully annotated images. This dataset was split into training (85% - 1168 images) and validation (15% - 206 images) sets. As preprocessing steps, each GFP max projection image was normalized and resized from 1024x1024 to 512x512. All ground-truth (GT) embryo instances were detected (true positives) with a Jaccard similarity score > 0.75 (Supplemental Figure 2a). Instances of false positives, where embryos that were not annotated as GT were detected, occurred in 7 (2.83%) predicted instances. Post-processing for each embryo included predicted label resizing back to the original size and creating a cropped individual embryo image (40 pixels padding) (Supplemental Figure 2a). Analysis scripts can be found here: https://github.com/PreibischLab/nd2totif-maskembryos-stagebin-pipeline/blob/master/2_stardist_pred <u>ict.py</u>

### Staging of embryos

Classification of different embryos into selected age bins was done by training an autoencoder-based classifier for stage prediction (Figure 2). The developmental bins used for staging (1-4, 5-30, 31-99, 100-149, 150-534, 535-558) were a compromise, selected based on biological meaning and classification prediction capability. A validation set comprising 250 embryos was employed to validate the autoencoder training. The validation of the classifier results was done using a class-stratified leave-one-out approach. For preprocessing of all images (training set and full dataset), 3D DAPI channel images were first masked for each embryo, then the 21 central slices were extracted, and pixel intensities were normalized using only non-zero values. Then, slightly modified copies were generated using small shifts, flips, rotations, shear, and brightness changes (augmentation). From those copies, 750 2D tiles of size 64x64 were extracted from random image regions for each embryo. All tiles of the embryo were used for training and inference.

First, an autoencoder was trained in an auto-associative, self-supervised manner to reconstruct its input, enabling it to learn a latent representation of the images. The autoencoder utilized mean squared error as the loss function and employed the Adam optimizer for training. In the subsequent step, the pre-trained encoder was integrated with a classifier designed to predict embryo stages, where the classification is achieved through a softmax output layer corresponding to the stage bins. This new network was initially trained with frozen encoder weights and then finally tuned by training the full network. For the classifier, cross-entropy was used as the loss function, and the Adam optimizer was used for optimization. A stage was predicted for each 64x64 tile of an embryo image, and the final stage for each embryo was determined by a majority vote of all tiles.

A training dataset was manually annotated using a custom-written ImageJ macro. We annotated ∼100 embryos with this approach, but we faced a strong class imbalance due to the initial random sampling of embryos. Therefore, an active learning approach (human in the loop) was utilized. Embryos for additional manual annotation were chosen based on low certainty in the classifier’s initial predictions, focusing on classes lacking sufficient examples. This method allowed us to move away from random sampling in subsequent iterations, strategically annotating stages with fewer examples to balance the distribution across classes (Figure 2B).

Scripts can be found here:

https://github.com/PreibischLab/nd2totif-maskembryos-stagebin-pipeline/blob/master/4_stage_predict ion.py

### Transcription detection

Detection of single RNA spots in 3D was performed using the Fiji plugin RS-FISH [51]. For this, images were preprocessed by subtracting a duplicated and median filtered (sigma = 19) image from the raw image to increase single spots and smooth background signals (Supplemental Figure 2B). Spot detection was performed using RS-FISH [51] with the following detection settings: -i0 0, -i1 65535, -a 0.650, -r 1, -s 1.09, -t 0.001, -sr 2, -ir 0.3, -e 0.5116, -it 33000, -bg 0. Spots were filtered using binary masks to exclude spots found from neighboring embryos using the Mask filtering option in RS-FISH with masks created using Stardist as described above. Computation was performed in parallel on the cluster and using AWS (Amazon Web Services). To correct for z-dependent signal loss, a quadratic function was fitted to the detected spots and used to correct the spot intensity throughout the embryo. A gamma function was fitted to the histogram of all found detections, and the maximum of the curve was set to 1 to normalize intensity detection between different embryos in order to allow quantification of transcript numbers (Figure 2B).

### Plotting and statistical analysis

Detected spots were plotted using custom-written scripts in R using ggplot2 and ggpubr. To test if mutant and wild-type conditions had significantly different RNA levels or intensities, a Welch’s unequal variances t-test was used in R. Significant P-values were indicated in the plots as n.s. (P > 0.05), * (P ≤ 0.05), ** (P ≤ 0.01), *** (P ≤ 0.001) and **** (P ≤ 0.0001). If one of the samples had too few points, no P-value was calculated.

### mRNA sequencing and analysis

mRNA-seq and data analysis from the deletion strain ERC41 were performed using methods to generate and analyze published data from wild-type embryos [64]. mRNA-seq data is deposited to GEO under series number GSE265989 with relevant information provided in Supplemental File 1.

## Acknowledgments

Research in this manuscript and SE was supported by the National Institute of General Medical Sciences of the National Institutes of Health under award number R35 GM130311. L. B. was supported by MDC-Berlin and the Joachim Herz Foundation (no. 850022), E. B. was supported by the HFSP grant RGP0021/2018-102 and MDC-Berlin, M. Z. was supported by EU H2020 Training grant 721890 ‘circRtrain’ and HHMI Janelia,. S. P. was supported by HFSP grant RGP0021/2018-102, EU H2020 Training grant 721890 ‘circRtrain,’ MDC-Berlin and HHMI Janelia. We thank Sarah Albritton and Haoyu Wang for their contribution to the mRNA-seq data and the Gencore at the NYU Center for Genomics and Systems Biology for sequencing and raw data processing. We additionally thank Andrew Woehler and the BIMSB advanced imaging facility for their support and maintenance of the microscope. We also thank the HHMI Janelia Open Science Software Initiative for maintenance and help with RS-FISH.

## Data availability

All raw smFISH images of *C. elegans* embryo, including the annotated data used for training, can be found here: https://data.janelia.org/3Mq4F. The RNA-seq data is deposited at Gene Expression Omnibus under series number GSE265989.

## Code availability

All custom code is either deposited or linked in this GitHub repository: https://github.com/ercanlab/2024_Breimann_et_al

## Supplemental Material

**Supplemental Figure 1.**
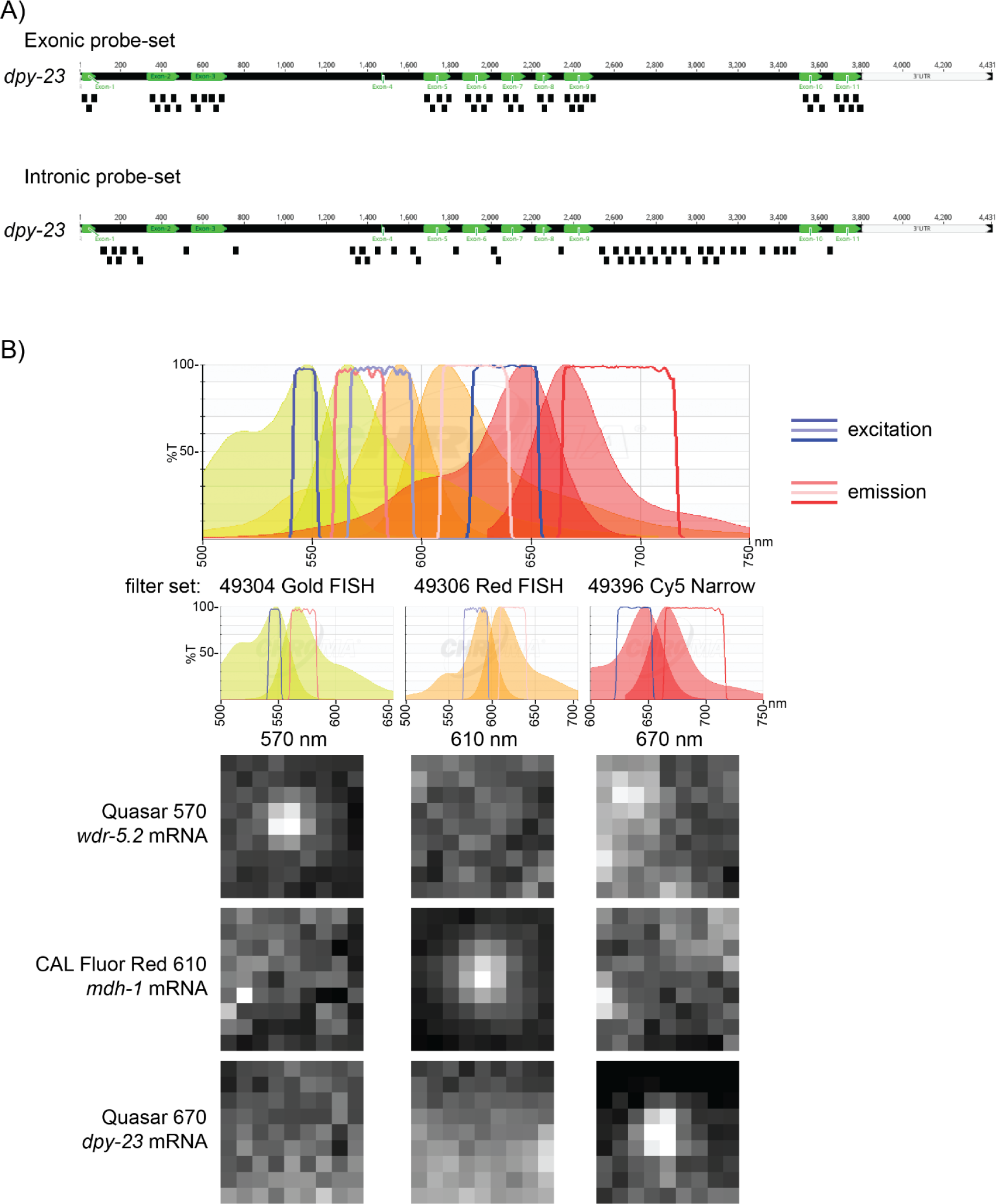
smFISH probe design. **A)** Example for exonic and intronic probes for DPY-23 mRNA**. B)** Curves of the fluorescent spectra of the three smFISH fluorophores (Quasar 570, CAL Fluor Red 610, and Quasar 670) and filter bandwidth for the three filters (Gold FISH, Red FISH, Cy5 Narrow) to specifically detect them. Spectra were created using the Chroma Spectra Viewer. Fluorescent images of smFISH spots for three different mRNAs with three different fluorophores. Images of the same spot in the different channels show the specificity of the signal and low bleed-through.

**Supplemental Figure 2.**
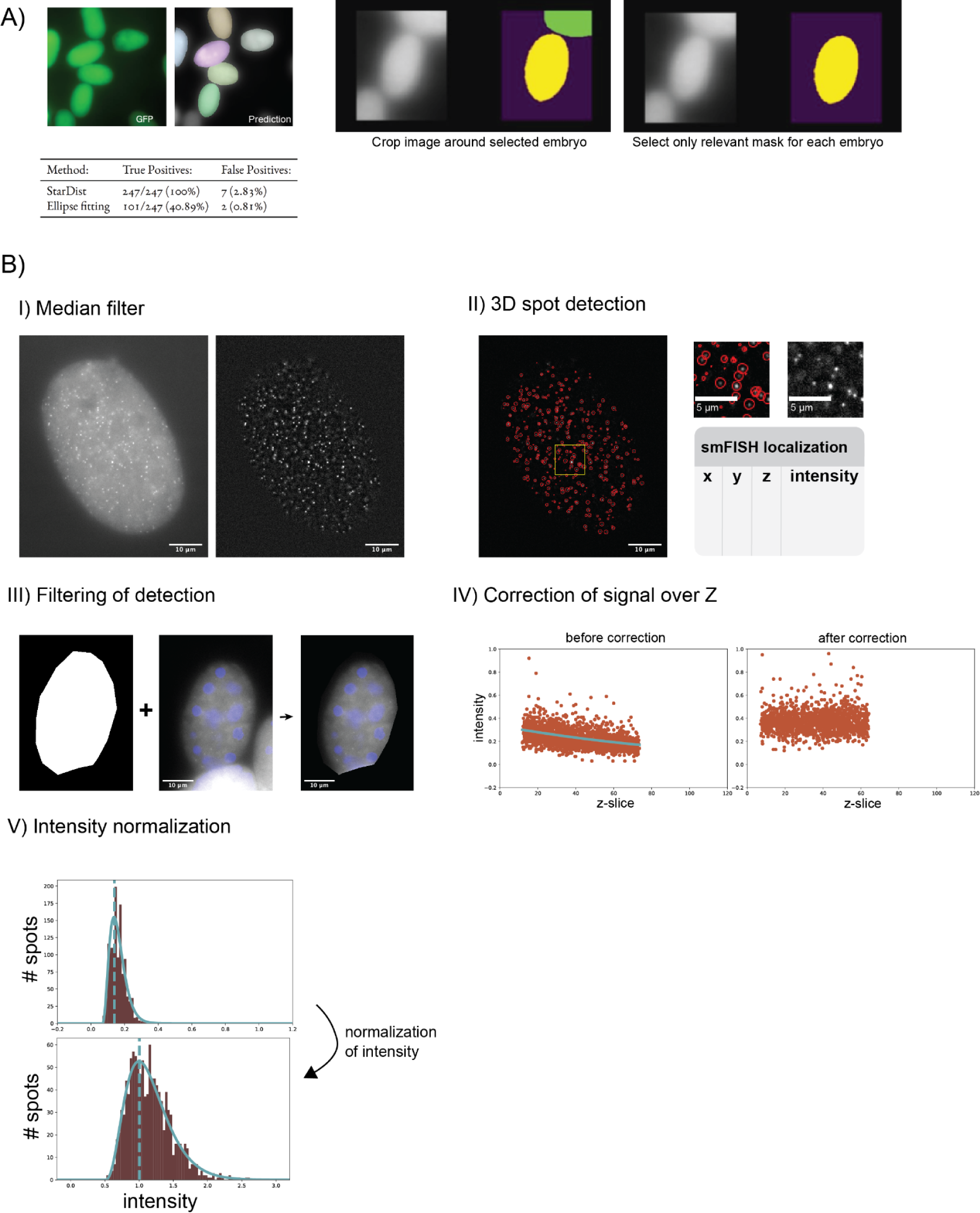
**A)** Instance segmentation using StarDist. The neural network was trained by the ground truth data created in the ellipsoid fitting. The network predicted masks on the 2D max projections of GFP images. Masks for single embryos are curated, and the original image is cropped around each segmented embryo. Jaccard similarity score was computed per instance in the validation set. Accepted correctly predicted instances were defined by a Jaccard score > 0.75. Using StarDist, all (100%) instances in the GT images were found. However, in rare cases (<3%), areas with low-quality embryos that are not desirable for analysis, as well as dirt, were detected. **B)** (I) The raw smFISH signal (left image) is filtered using a median filter (sigma =19) to enhance the spots for detection (right image). (II) Single mRNAs are detected in 3D using RS-FISH, which records a table of localizations and intensity. (III) Masks (left image) are created using Stardist, which are used to filter localizations (right image). (IV) The image intensity drops as a function of distance to the coverslip (z) (left plot). Through fitting a quadratic function, the intensities are corrected in z (right plot). (V) The intensity is normalized by fitting a gamma function, where the maximum is set to 1.

**Supplemental Figure 3.**
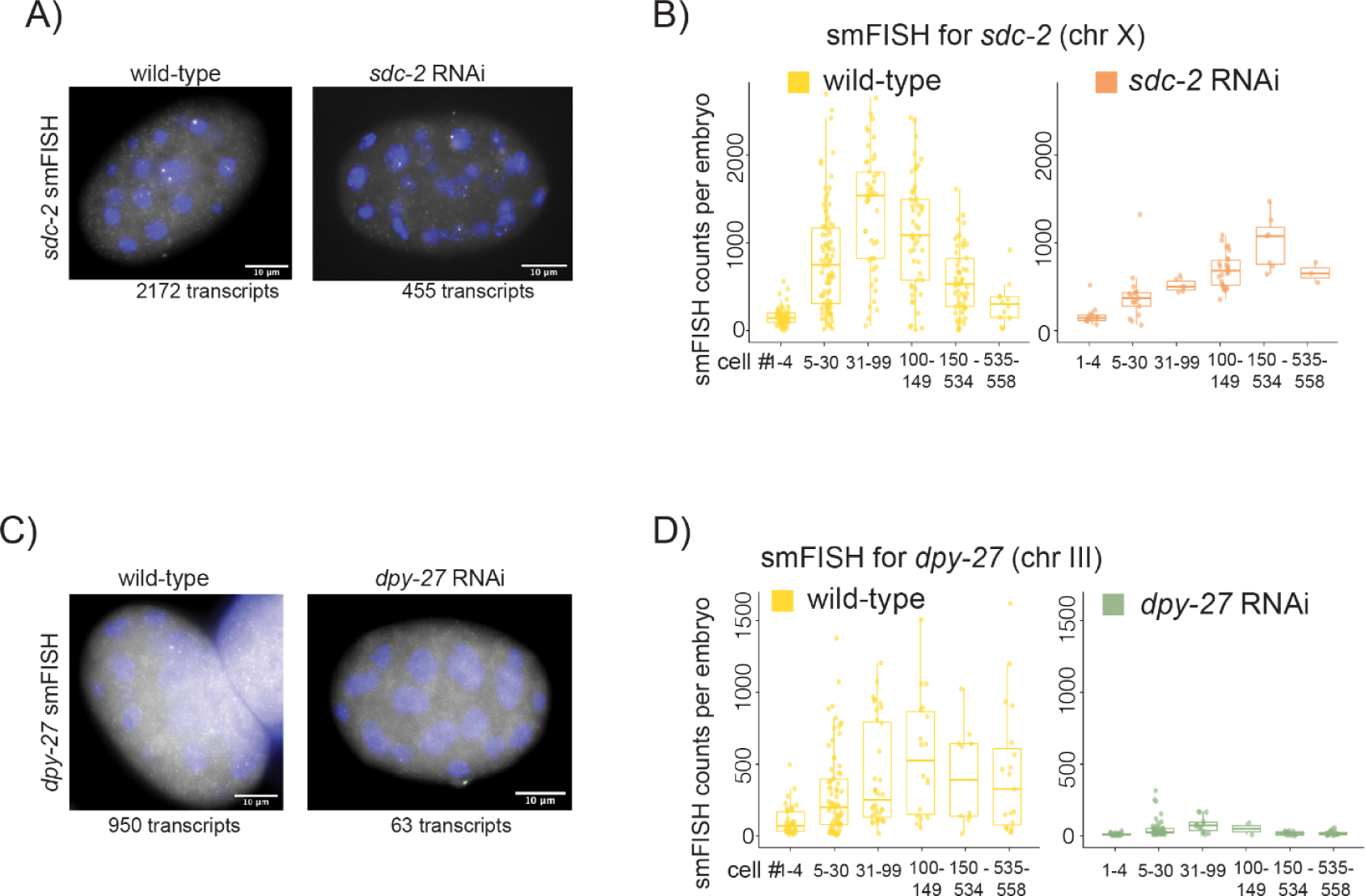
**A)** Example images of wild-type or *sdc-2* RNAi embryos labeled with *sdc-2* smFISH. The number of detected smFISH spots for each example is written under the image. **B)** Plots for smFISH detection of X chromosomal mRNA *sdc-2*. Each spot represents the total mRNA count for one embryo, plotted against the binned nuclei number for wild type (yellow) and *sdc-2* RNAi (orange). **C)** Example images of wild-type or *dpy-27* RNAi embryos labeled with *dpy-27* smFISH. The number of detected smFISH spots for each example is written under the image. **D)** Plots for the autosomal mRNA *dpy-27* in wild-type (yellow) or *dpy-27* RNAi (green) conditions.

**Supplemental Figure 4.**
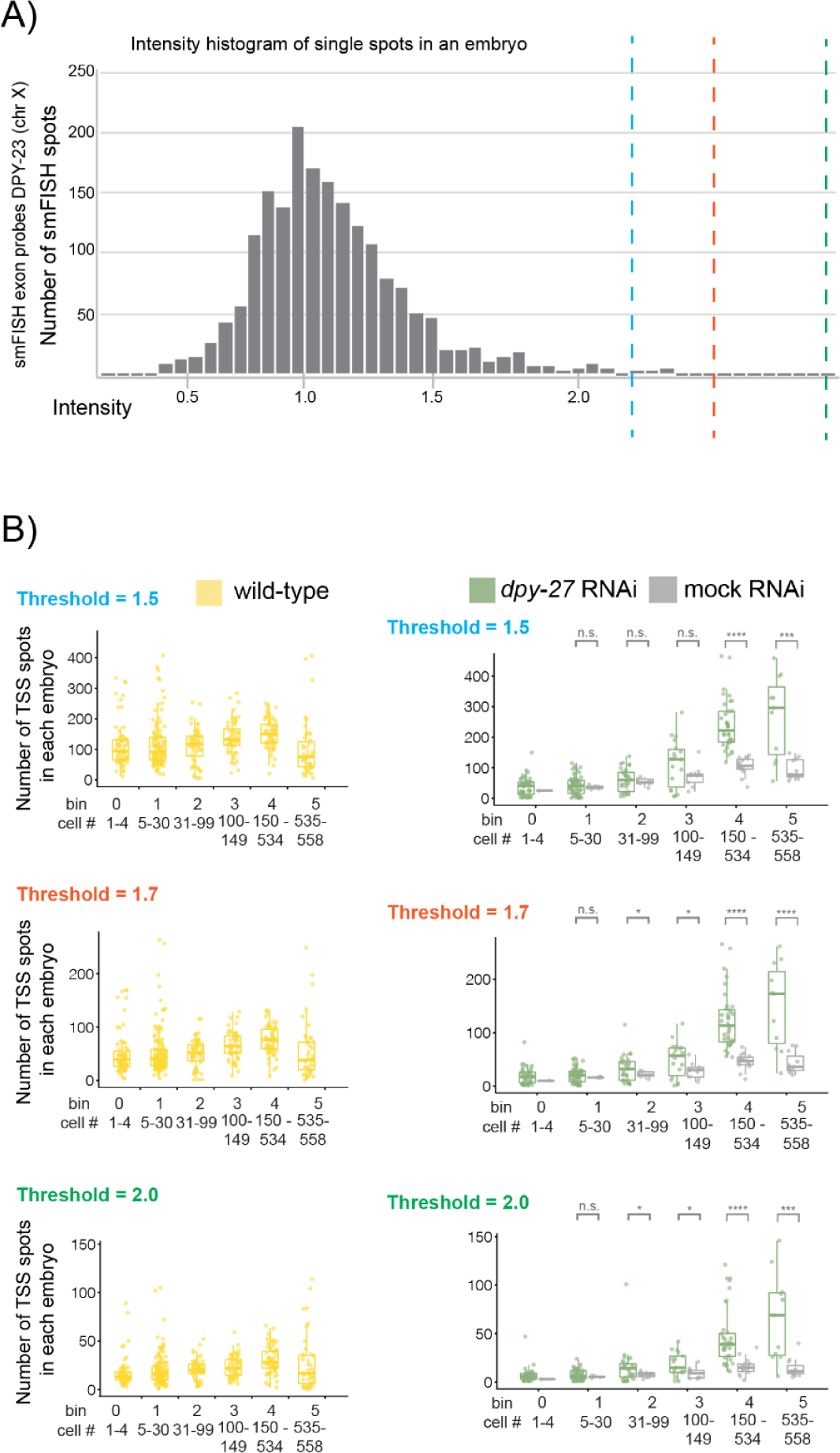
Analysis of threshold selection for transcription start site. **A)** Histogram of the normalized exon intensities of all detected spots for one embryo. Different cut-offs for detection of high-intensity spots were selected at 1.5 (blue), 1.7 (red), and 2.0 (green). **B)** Relative change of actively transcribing promoters for X chromosomal genes for wild-type (yellow), *dpy-27* RNAi (green), and mock RNAi (gray) conditions. Three different thresholds were selected to classify transcription sites in the dataset. Significant P-values from Welch’s unequal variances t-test are noted above the wild-type vs. knock-down condition for each time-point. P-values are denoted as n.s. (P > 0.05), * (P ≤ 0.05), ** (P ≤ 0.01), *** (P ≤ 0.001) and **** (P ≤ 0.0001). If one of the samples had less than two points, no P-value was calculated.

**Supplemental Figure 5.**
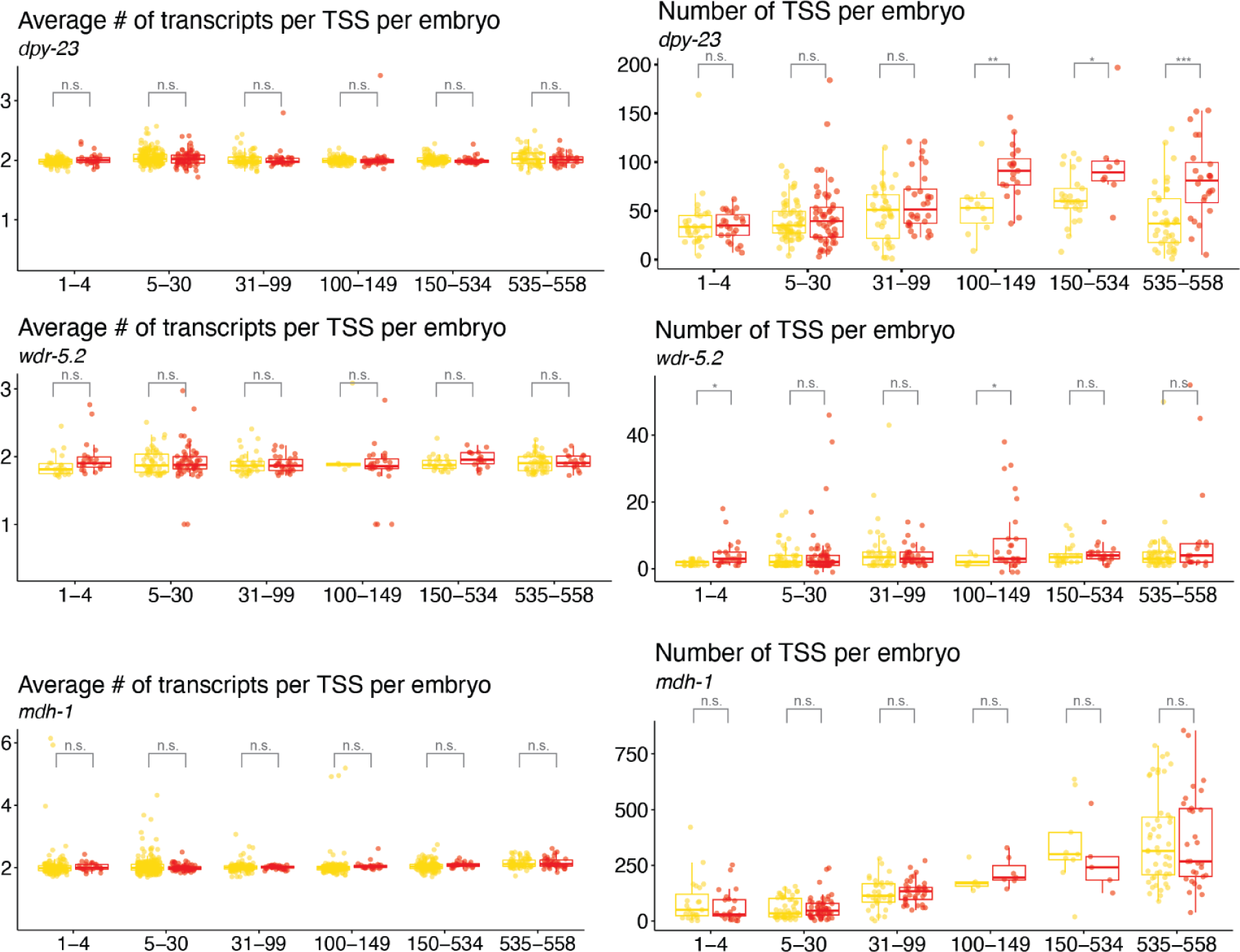
Analysis of transcription site parameters in *rex-1* deletion. Relative transcription parameter (average number of transcripts per transcription site per embryo and the number of transcription spots per embryo) changes for the X chromosomal genes *dpy-23* and *wdr-5.2*, as well as autosomal gene *mdh-1* for wild-type (yellow) and rex-1 deletion (red) conditions. Data are from embryos probed for the three genes simultaneously Significant P-values from Welch’s unequal variances t-test are noted above the wild-type vs. knock-down condition for each time-point. P-values are denoted as n.s. (P > 0.05), * (P ≤ 0.05), ** (P ≤ 0.01), *** (P ≤ 0.001) and **** (P ≤ 0.0001).

### Information on the imaging data

The complete raw imaging data can be found at https://data.janelia.org/3Mq4F. Images are split into subfolders of genotypes N2 (wild-type, CGC), CB428 (dpy-21 (e428), SEA-12 (ERC41 (ers8[delX:4394846-4396180]))) or RNAi treatment (sdc-2 (C35C5.1), dpy-27 (R13G10.1), ama-1 (F36A4.7), ev (empty-vector), rluc (Renilla luciferase)). Additionally, the embryo masks are provided in the file mask.zip. The manual annotation of the embryo stage and the image tiles used in the autoencoder’s training can be found in the manual_count.txt and tiles.zip, respectively.

**Table 1.** The number of probes and fluorophore of each probe-set.

| gene | Chr | transcript<br>length<br>(nt) | N2<br>FPKM |  | Vector<br>RNAi<br>FPKM | RNAi<br>DPY-27<br>FPKM | DPY-21 KO FPKM |  | Mature RNA |  | Nascent RNA |  |
| --- | --- | --- | --- | --- | --- | --- | --- | --- | --- | --- | --- | --- |
|  |  |  | Early<br>embryo | Comma<br>embryo | Mixed<br>embryo | Mixed<br>embryo | Early<br>embryo | Mixed<br>embryo | # of<br>probes | Fl | # of<br>probes | Fl |
| *dpy-23* | X | 1980 | 68,64 | 76,93 | 61,98 | 79,23 | 66,81 | 89,03 | 48 | Q670 | 48 | Q570 |
| *sdc-2* | X | 9247 | 27,79 | 10,26 | 14,65 | 21,54 | 25,26 | 14,21 | 48 | Q670 | 48 | Q570 |
| *wdr-5.2* | X | 2040 | 23,06 | 34,85 | 29,67 | 39,67 | 13,66 | 30,47 | 48 | Q570 | - | - |
| *rab-6.2* | X | 913 | 53,11 | 206,39 | 178,36 | 193,41 | 42,31 | 170,27 | 41 | Q670 | 46 | Q570 |
| *mdh-1* | V | 1149 | 123,26 | 588,64 | 634,14 | 599,16 | 121,53 | 601,70 | 41 | C610 | - | - |
| *ama-1* | IV | 6020 | 271,49 | 92,02 | 92,88 | 99,41 | 255,66 | 104,29 | 48 | C610 | 48 | Q570 |
| *dpy-27* | III | 4677 | 58,82 | 77,45 | 69,78 | 26,95 | 80,74 | 56,82 | 48 | C610 | - | - |

**Table 2.**
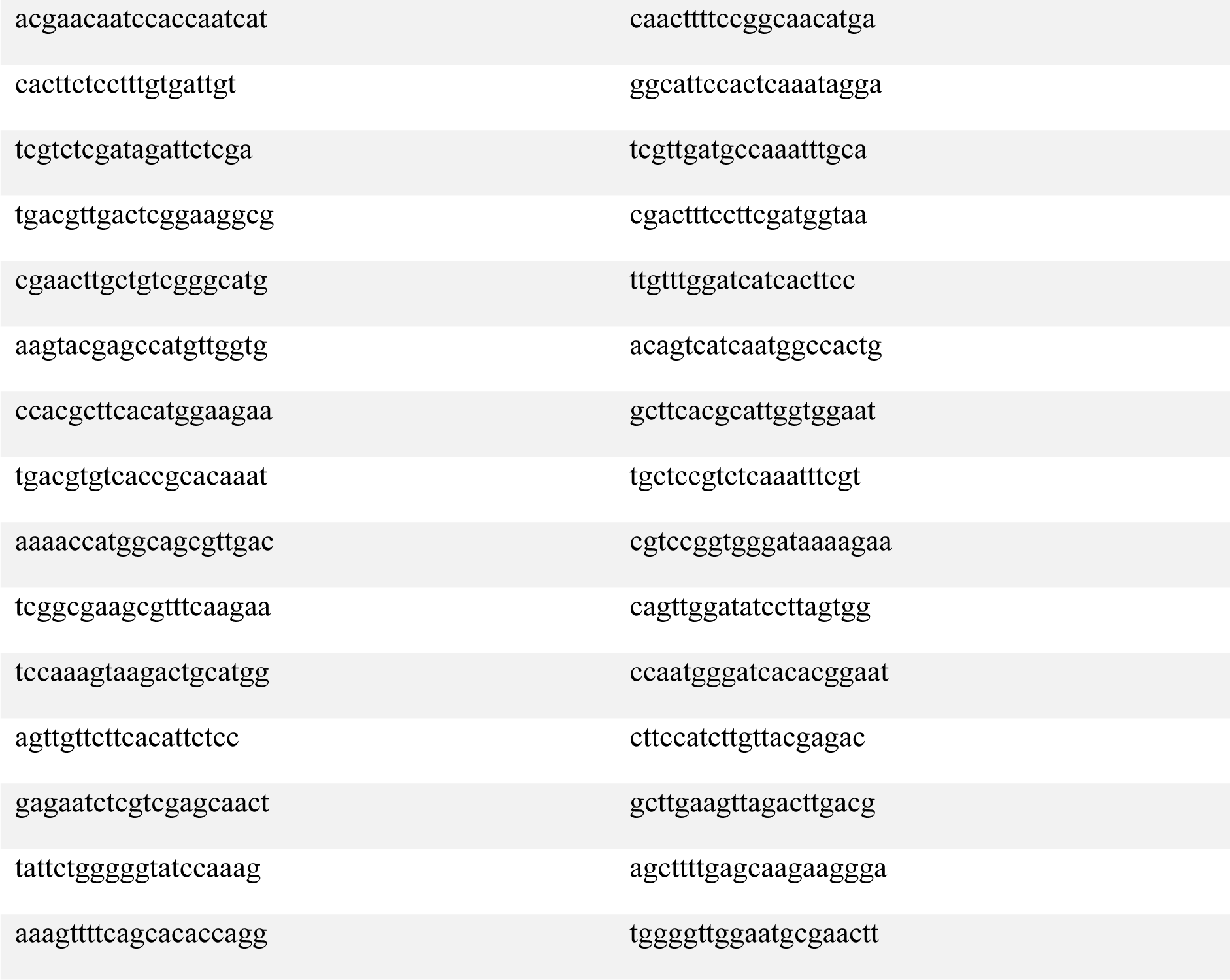
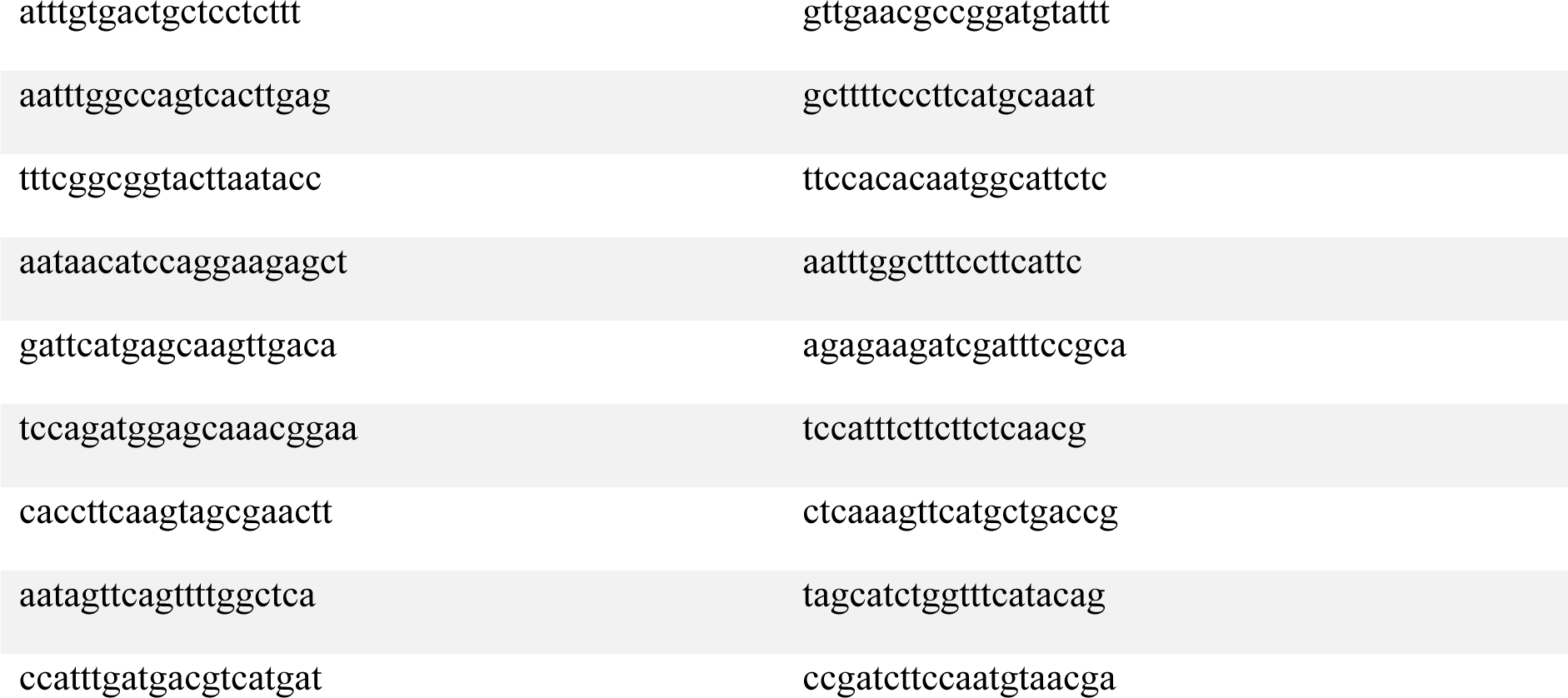
DPY-23 exonic probes.

**Table 3.**
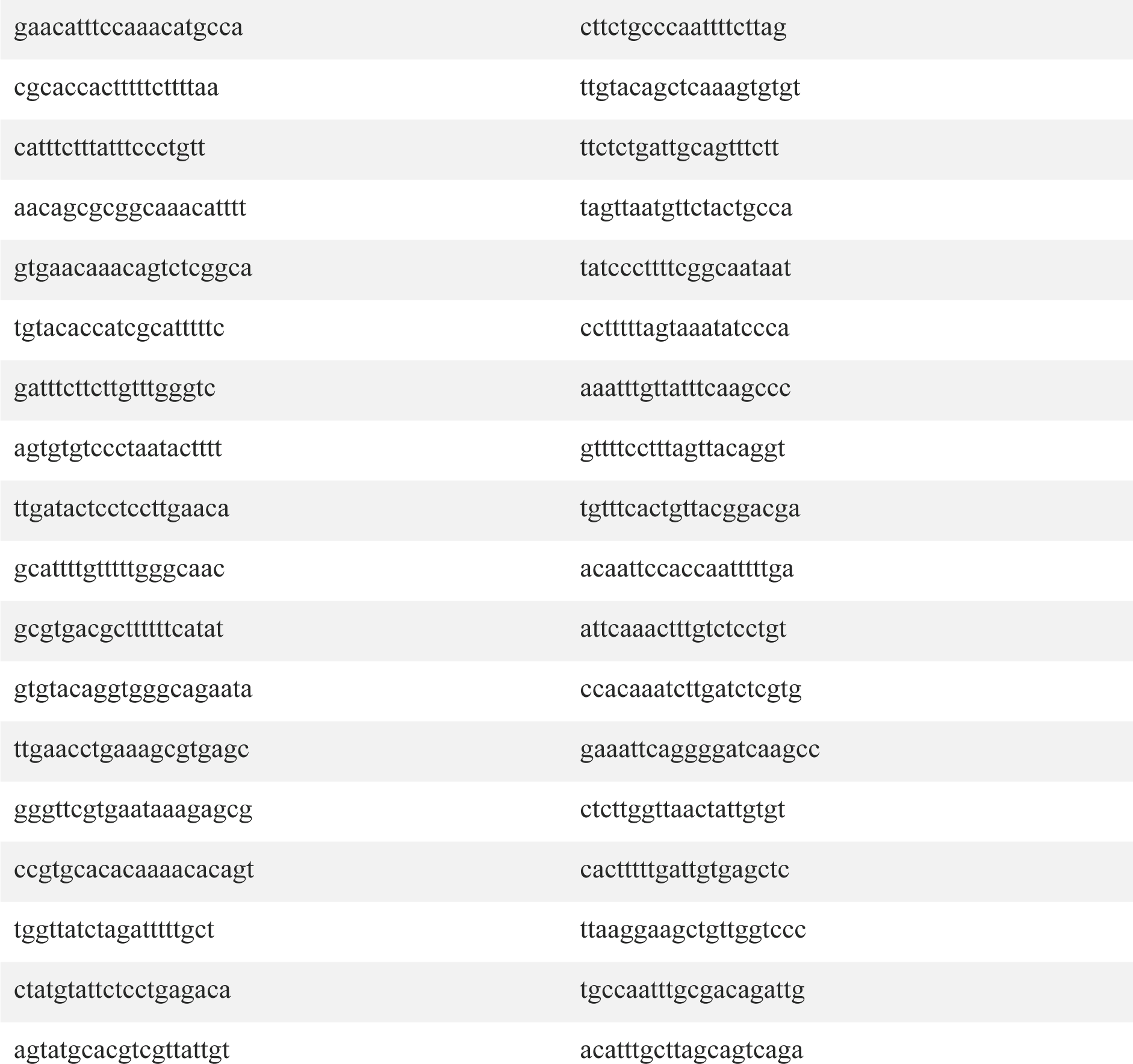
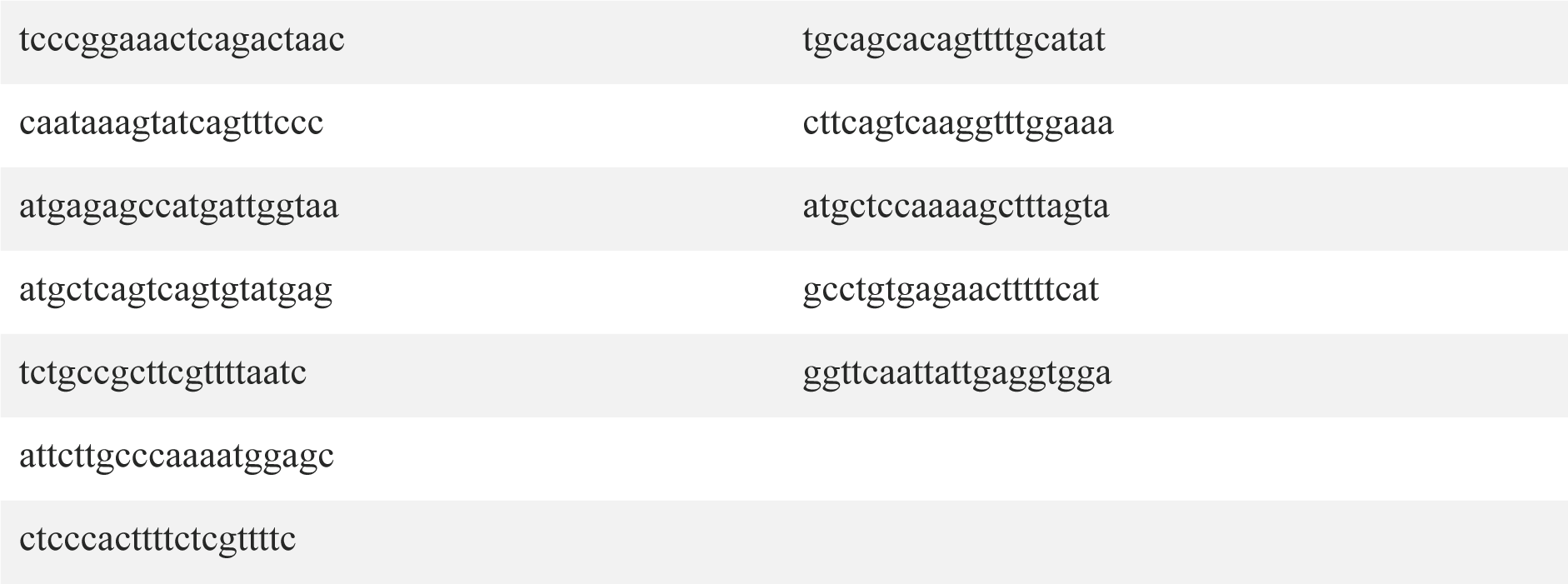
DPY-23 intronic probes.

**Table 4.**
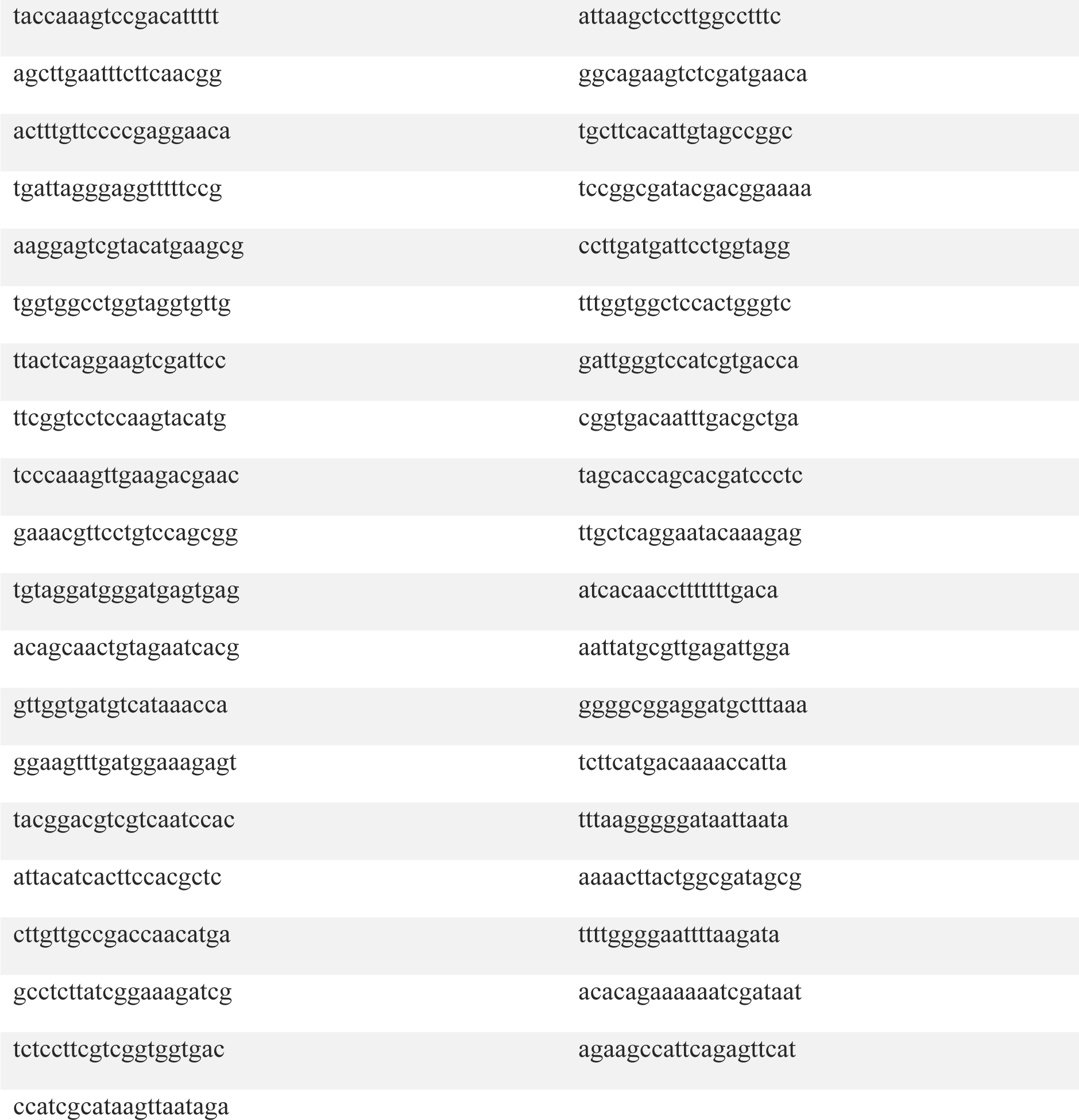
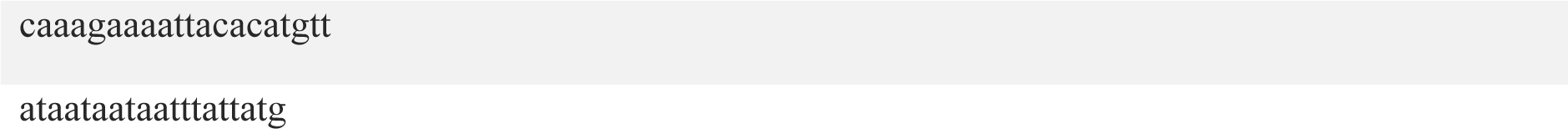
RAB-6.2 exonic probes.

**Table 5.**
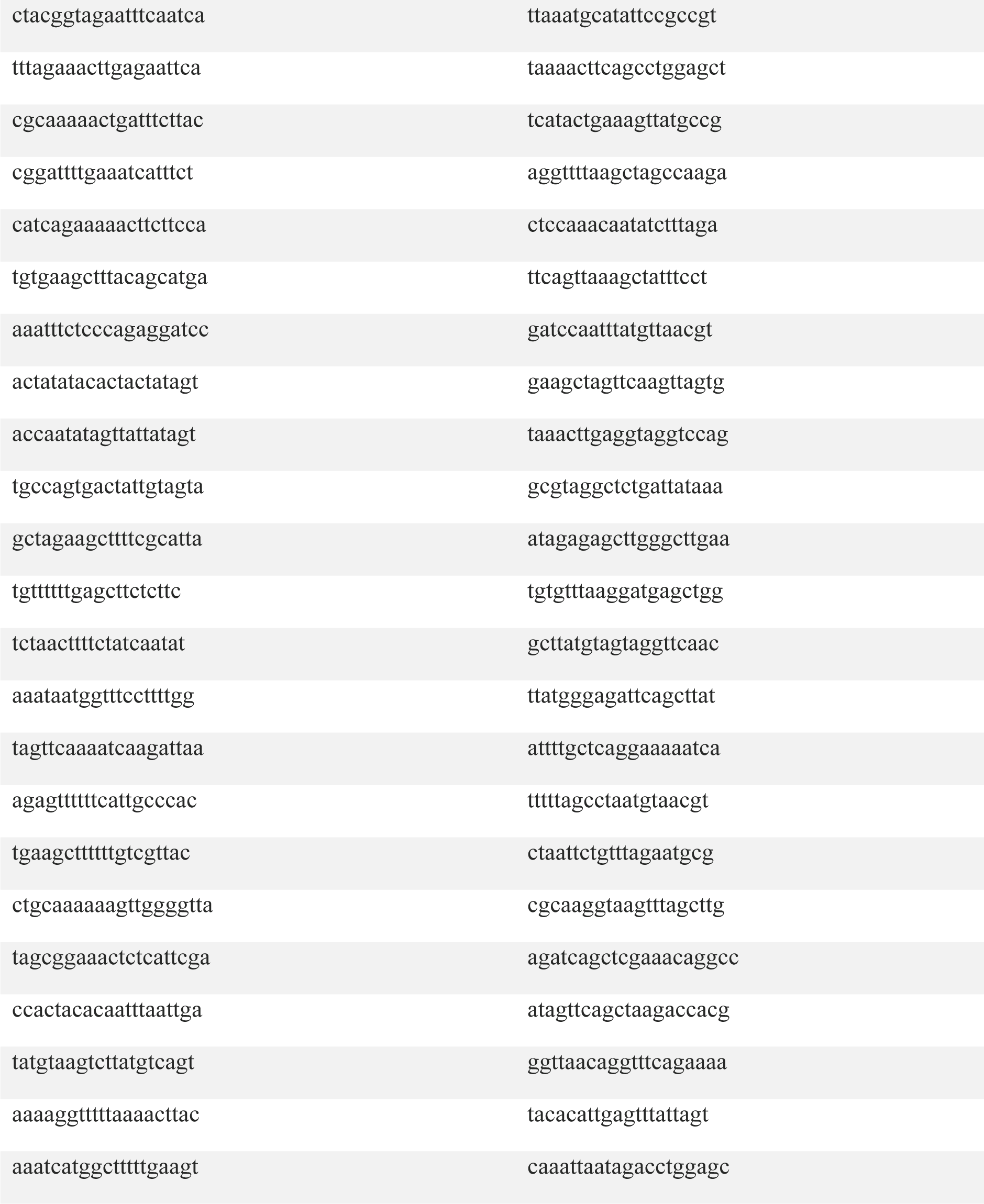
RAB-6.2 intronic probes.

**Table 6.**
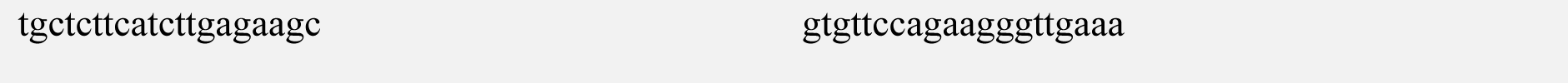
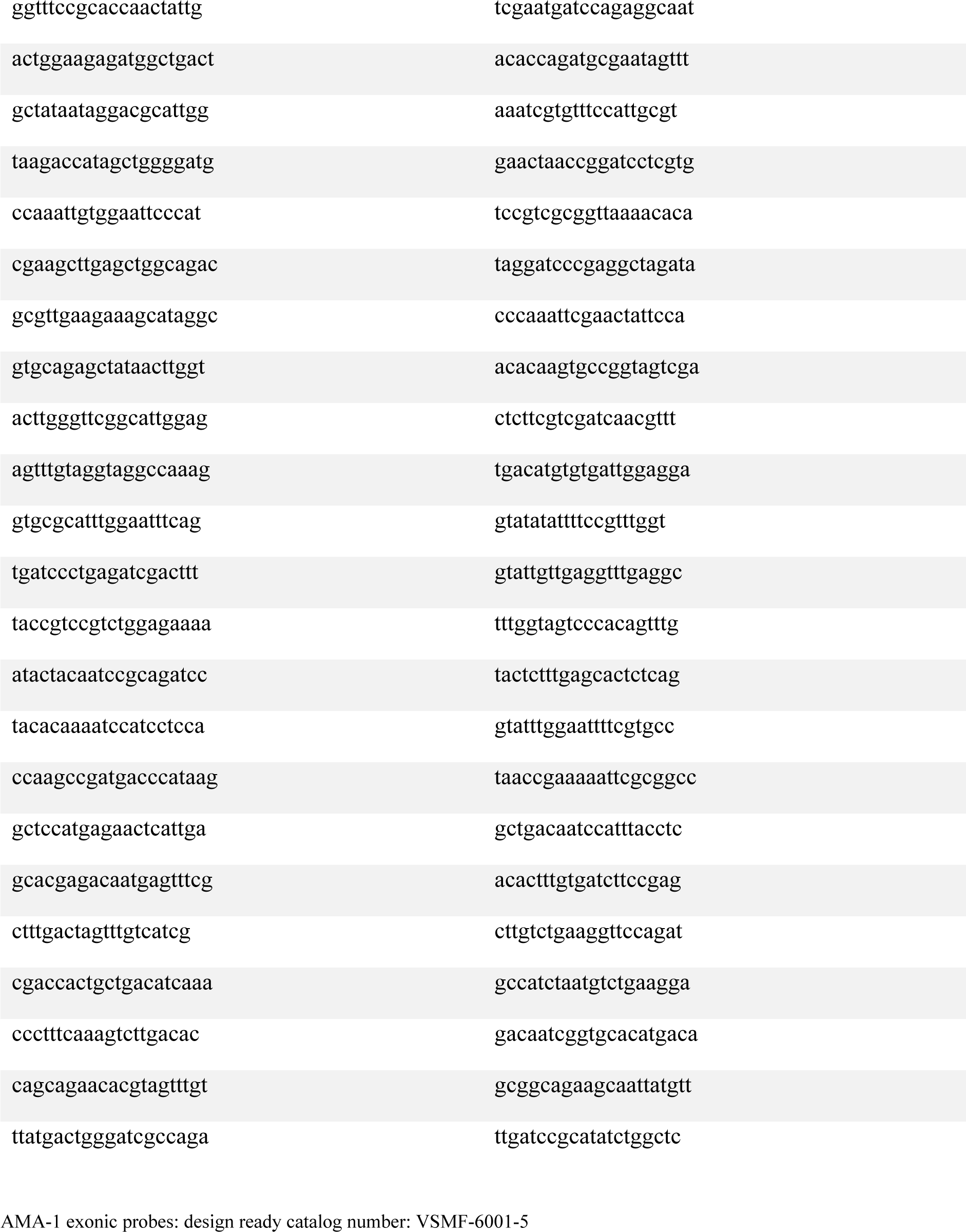
WDR-5.2 exonic probes.

**Table 7.**
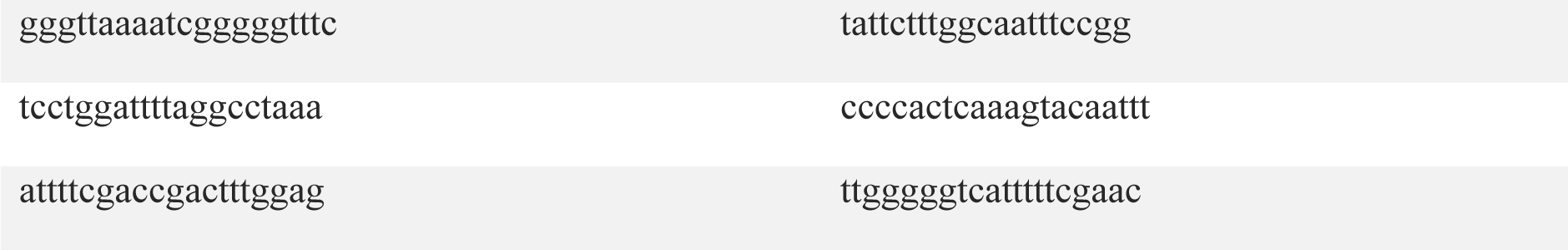
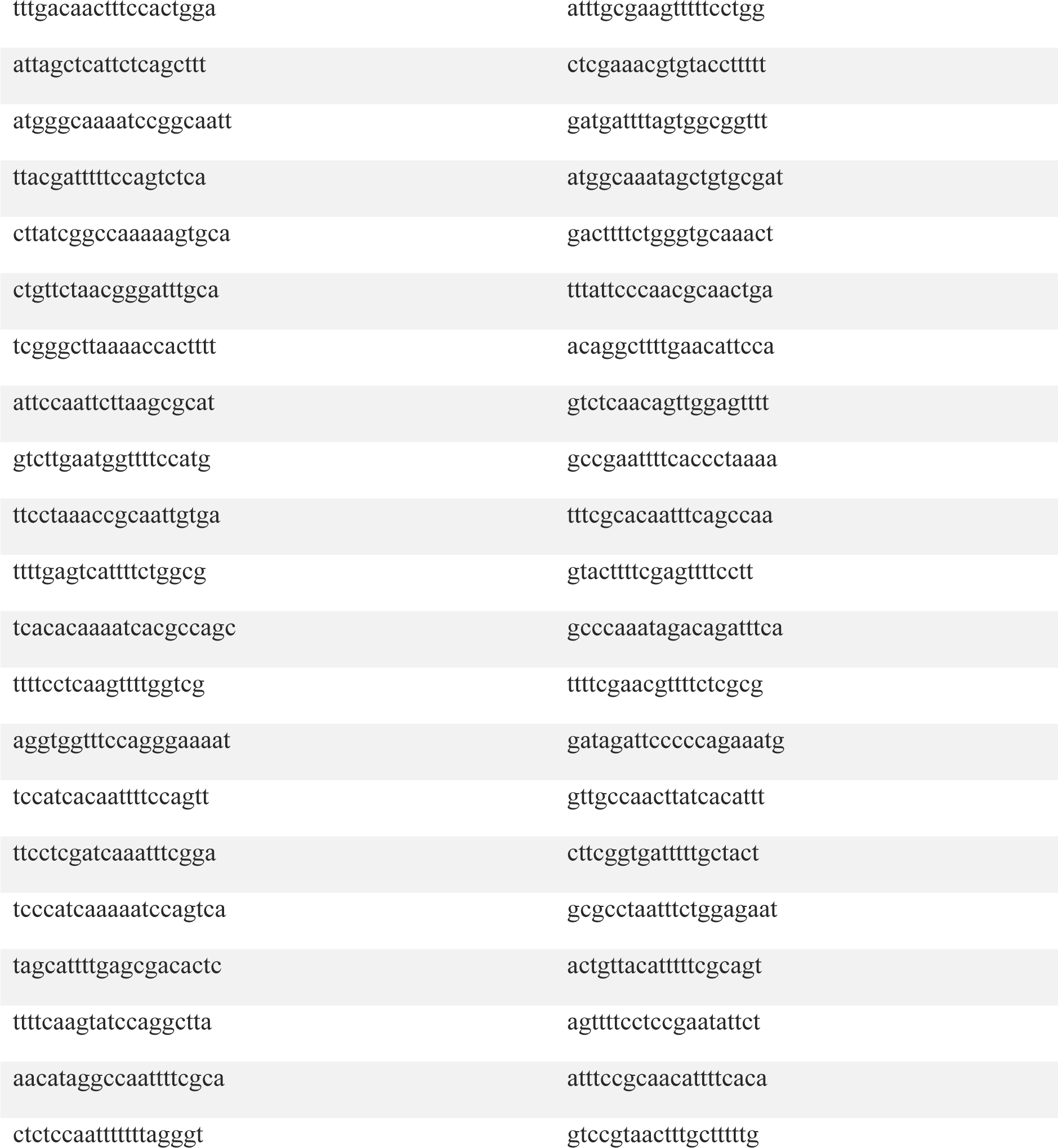
AMA-1 intronic probes.

**Table 8.**
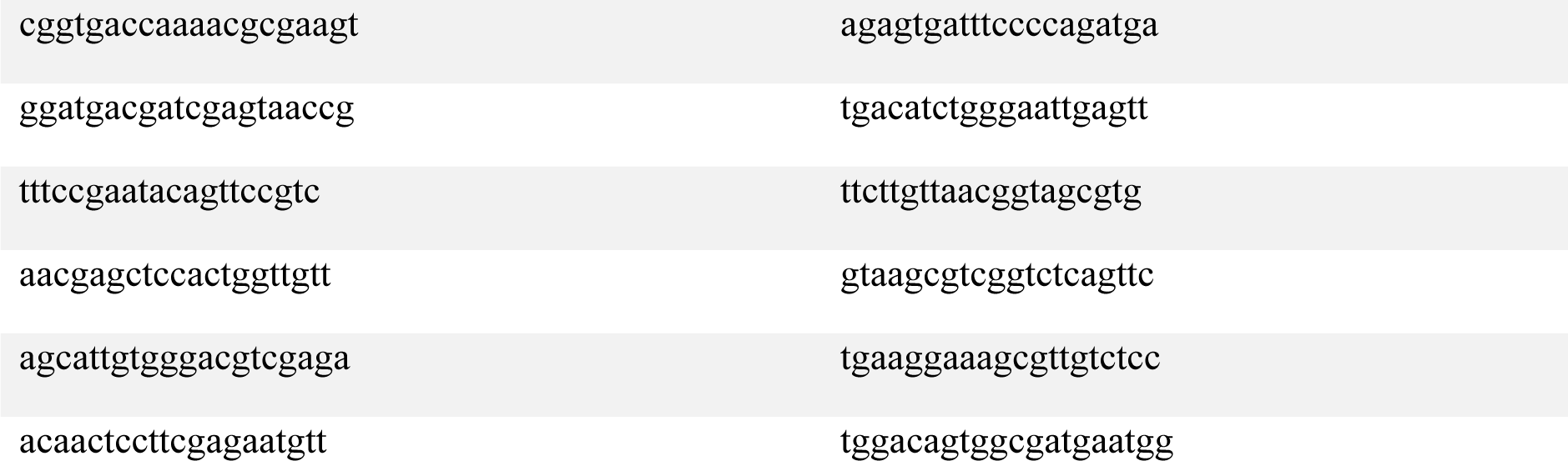
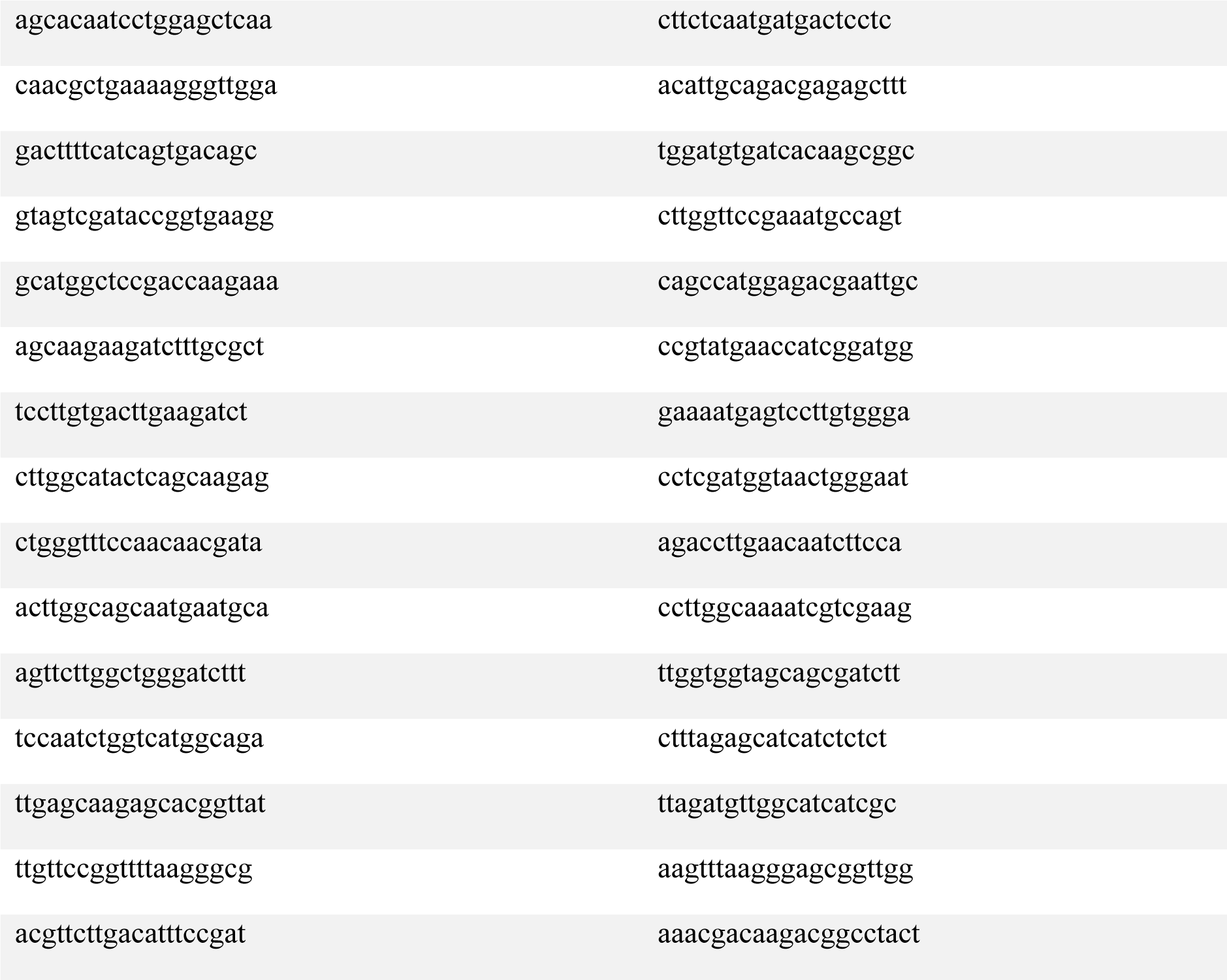
MDH-1 exonic probes.

**Table 9.**
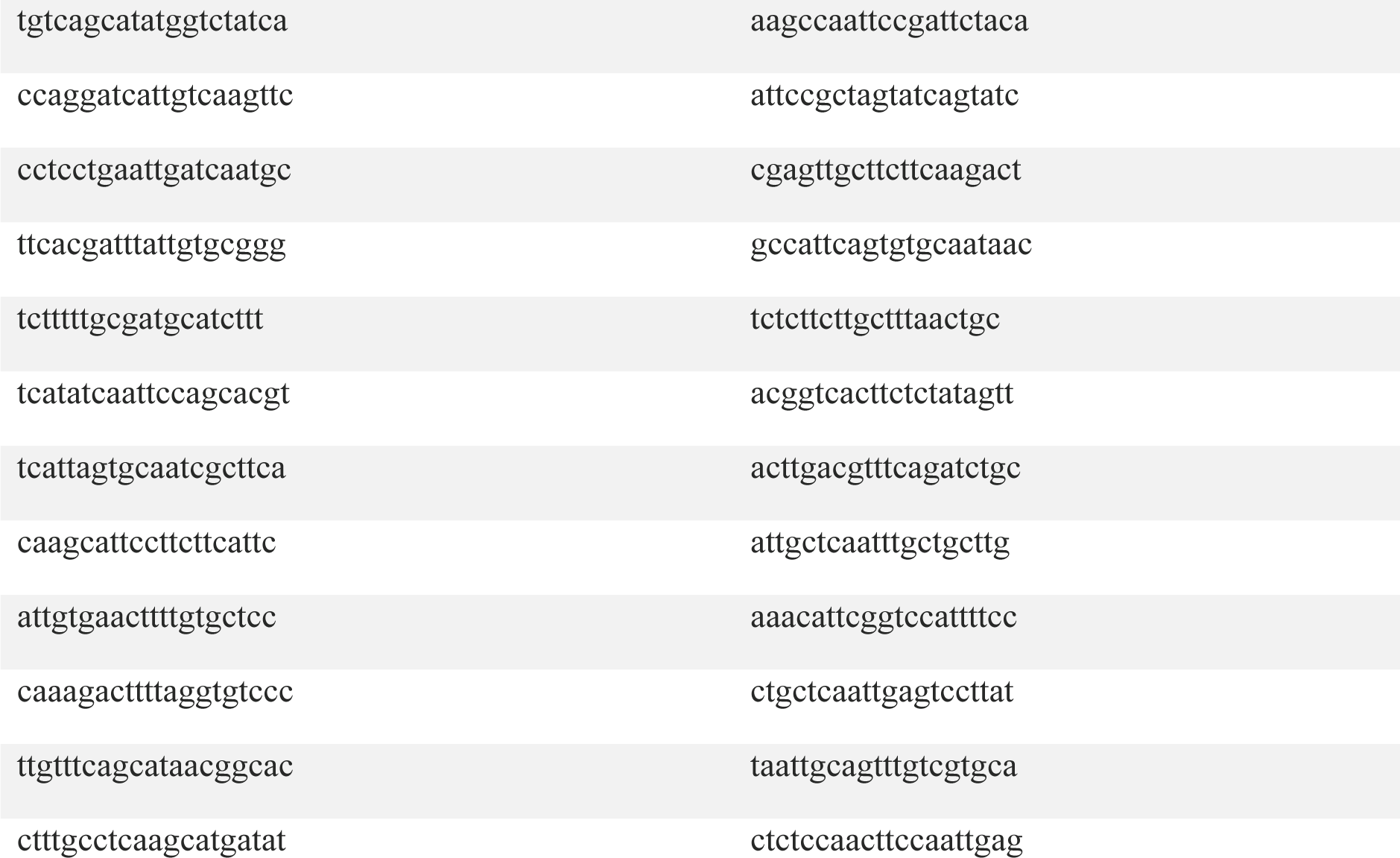
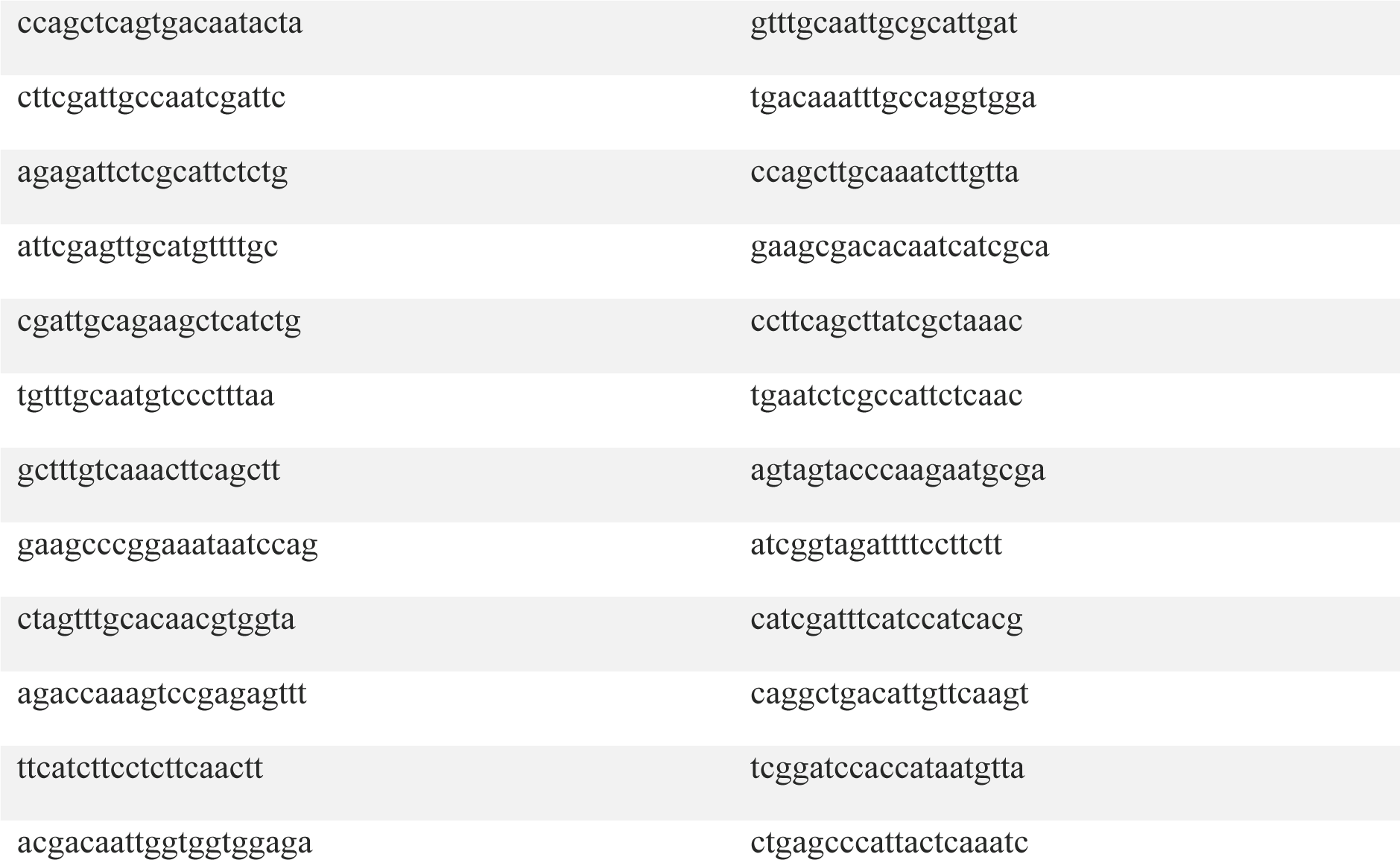
DPY-27 exonic probes.

**Table 10.**
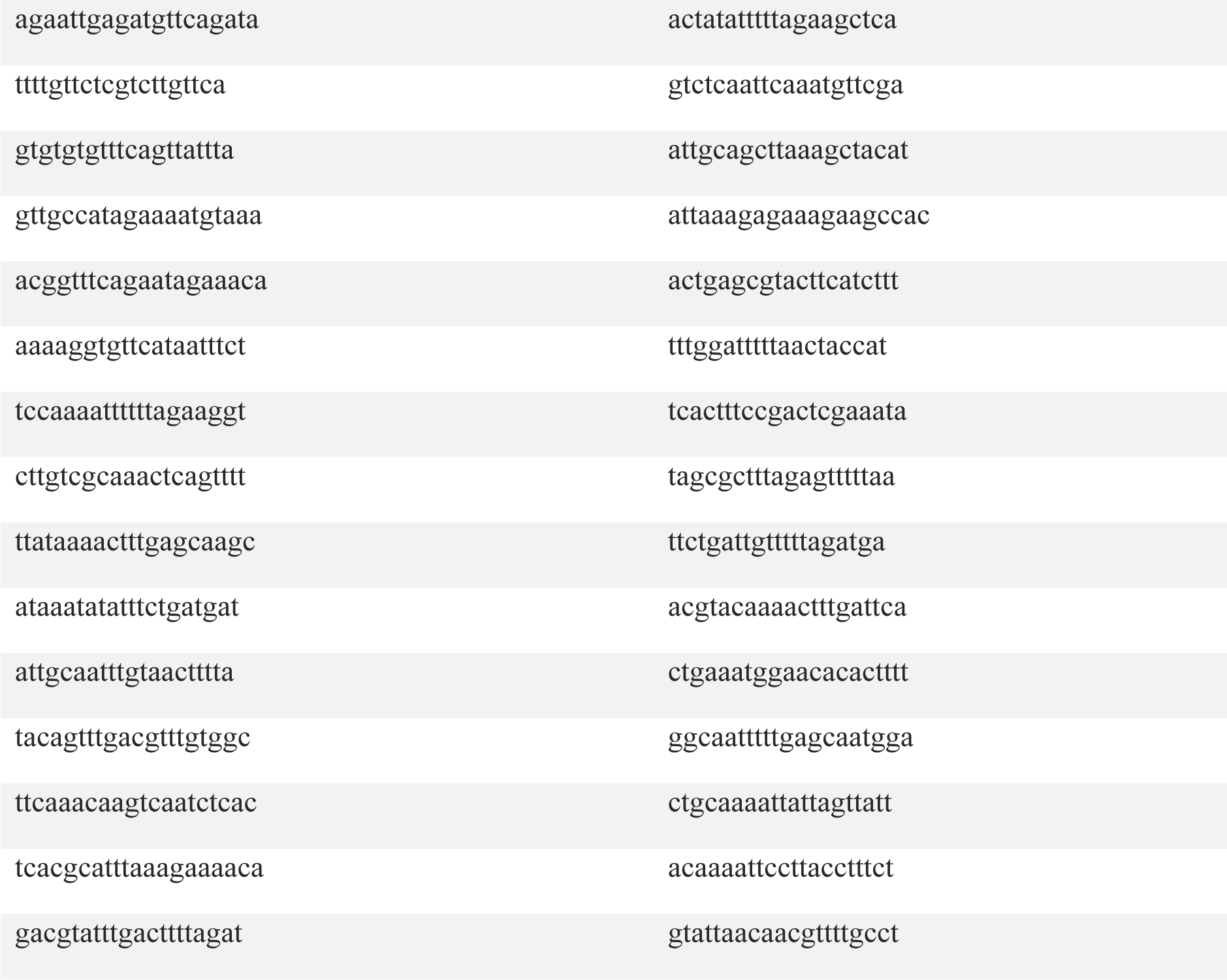
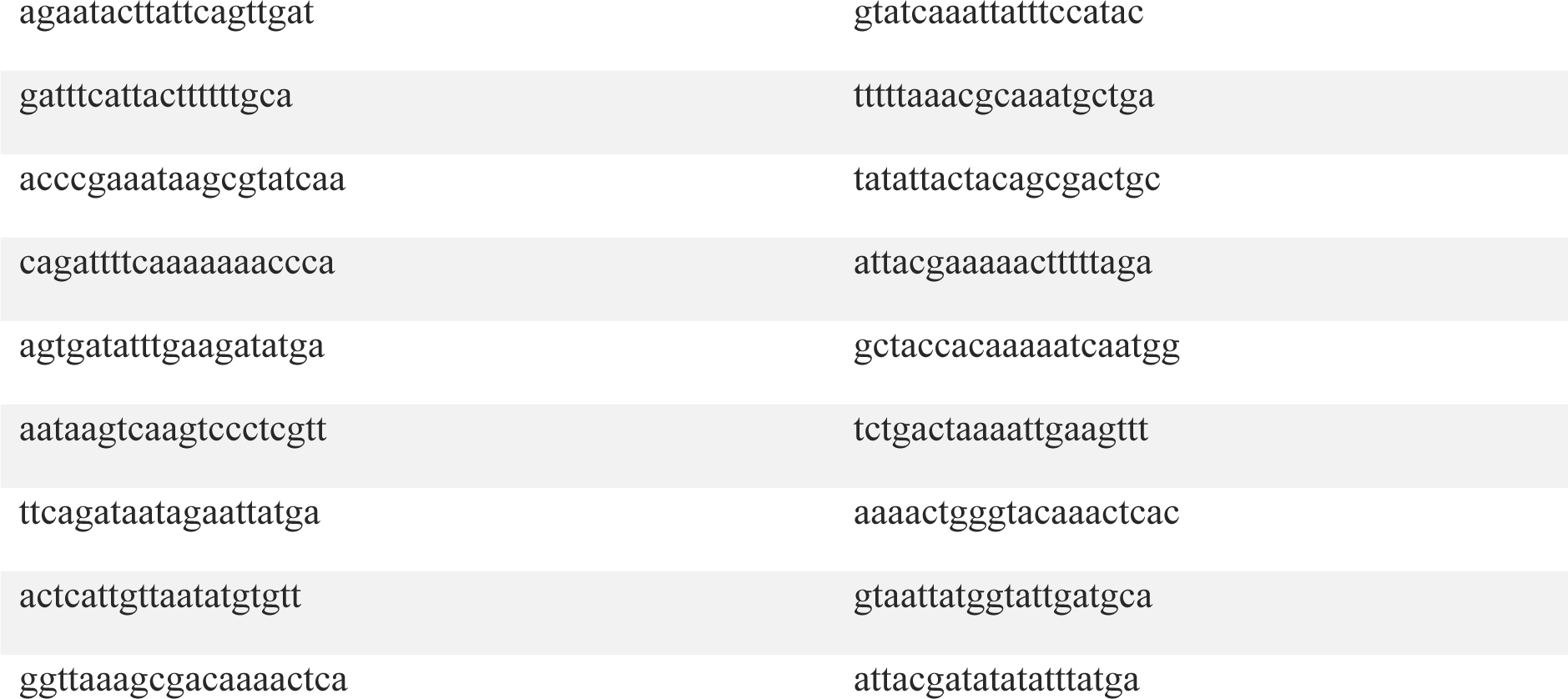
SDC-2 intronic probes.

**Table 11.**
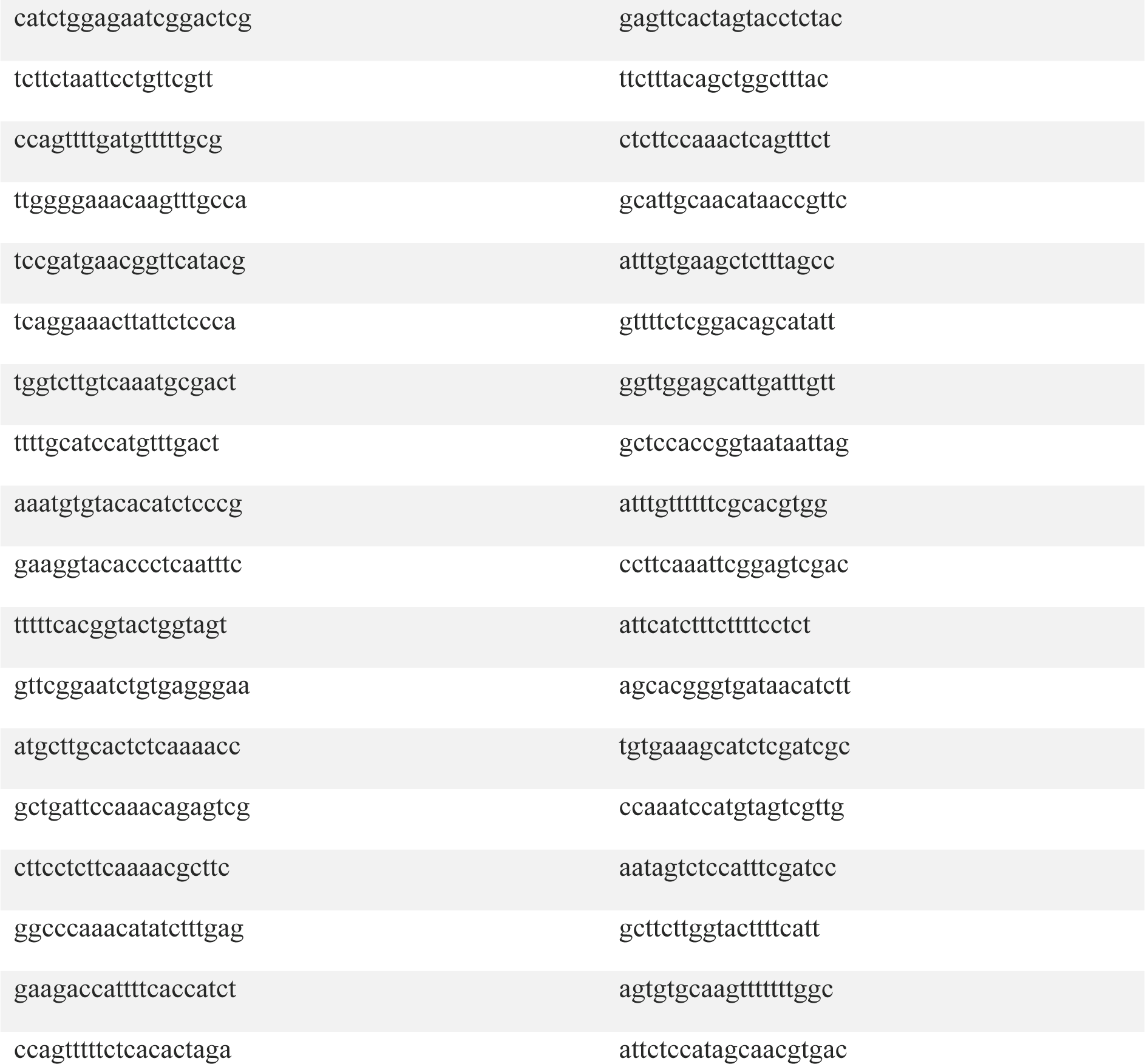
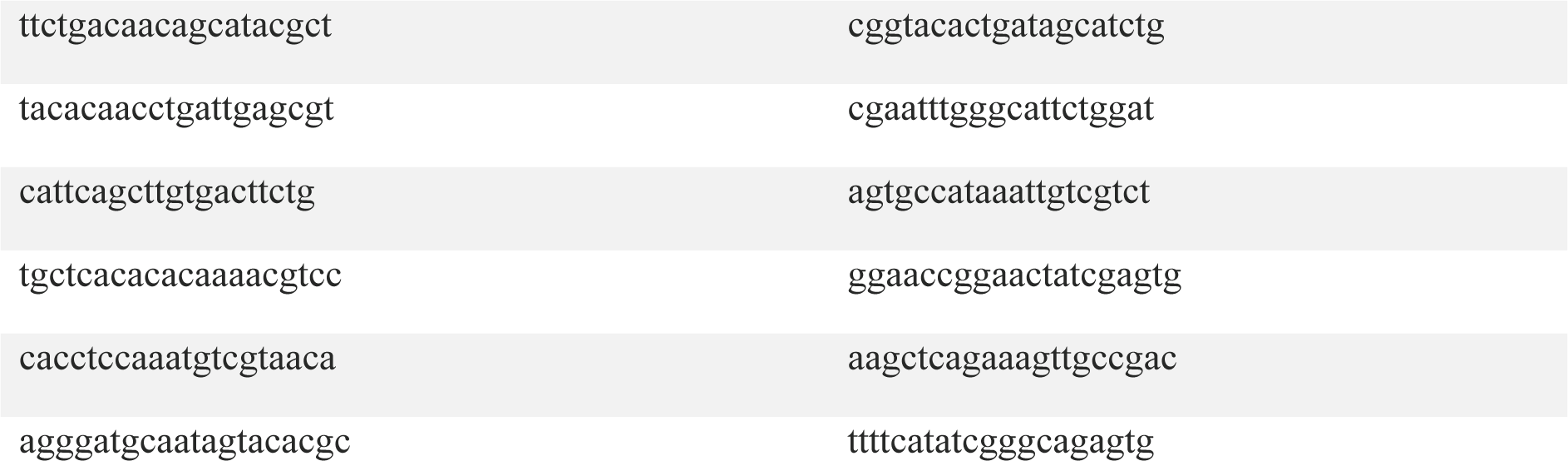
SDC-2 exonic probes.

